# Interaction of sortilin with apolipoprotein E3 enables neurons to use long-chain fatty acids as alternative metabolic fuel

**DOI:** 10.1101/2024.06.10.598173

**Authors:** Anna K Greda, Jemila P Gomes, Ewa Zurawska-Plaksej, Raphaela Fritsche-Guenther, Ina-Maria Rudolph, Narasimha S Telugu, Cagla Cömert, Jennifer Kirwan, Séverine Kunz, Michael Rothe, Sebastian Diecke, Peter Bross, Thomas E Willnow

**Affiliations:** Departments of Biomedicine, Aarhus University, Aarhus, Denmark; Molecular Cardiovascular Research and Technology Platforms for, Max-Delbrueck-Center for Molecular Medicine in the Helmholtz Association, Berlin, Germany; Department of Toxicology, Wroclaw Medical University, Wroclaw, Poland; Berlin Institute of Health at Charité – Universitätsmedizin Berlin, Metabolomics, Berlin, Germany; Pluripotent Stem Cells, Max-Delbrueck-Center for Molecular Medicine in the Helmholtz Association, Berlin, Germany; Departments of Clinical Medicine, Aarhus University, Aarhus, Denmark; Electron Microscopy, Max-Delbrueck-Center for Molecular Medicine in the Helmholtz Association, Berlin, Germany; Lipidomix GmbH, Berlin, Germany

## Abstract

Sortilin (*SORT1*) is a lipoprotein receptor that shows genome-wide association with hypercholesterolemia, explained by its ability to control hepatic output of lipoproteins. Remarkably, *SORT1* also shows genome-wide association with Alzheimer disease (AD) and frontotemporal lobe dementia, the most prevalent forms of age-related dementias. Yet, sortilin’s contribution to human brain lipid metabolism and health remains unclear. Using humanized mouse strains and iPSC-based cell models of brain lipid homeostasis, we document that sortilin mediates neuronal uptake of polyunsaturated fatty acids carried by apoE. Internalized lipids are converted into ligands for PPARα, inducing transcription profiles that enable neurons to use long-chain fatty acids as metabolic fuel. This pathway works with apoE3, but is lost with the AD risk factor apoE4, which disrupts sortilin’s endocytic activity. We document a role for the lipoprotein receptor sortilin in metabolic fuel choice in neurons, possibly crucial when supply with glucose is limited, as in the aging brain.

## INTRODUCTION

Sortilin is a 95 kDa type-1 transmembrane receptor expressed in various mammalian cell types, most notably in neurons of the central and peripheral nervous system and in hepatocytes in the liver^1–3^. Sortilin operates as an endocytic and intracellular sorting receptor, directing cargo between the cell surface and various compartments of biosynthetic and endocytic pathways (reviewed in ^4–6^). In the liver, it acts as a lipoprotein receptor controlling hepatic uptake but also release of cholesterol-rich lipoproteins, explaining genome-wide association of the encoding gene *SORT1* with plasma cholesterol levels and the risk of myocardial infarction^2,3,7–9^

Recently, we showed that sortilin also acts as a lipoprotein receptor in the brain, facilitating neuronal uptake of apolipoprotein (apo) E, the major carrier for lipids in brain interstitial fluids^10^. ApoE is secreted by astrocytes and microglia and delivers essential lipids to neurons (reviewed in ^11–13^). In mouse models, sortilin-dependent uptake of lipids works well with apoE3, the most common isoform of the lipid carrier in humans. However, this function of sortilin is disrupted when binding lipidated apoE4, as this apoE variant blocks sortilin recycling and ceases receptor-mediated ligand uptake into cells^14,15^. This observation is significant as *APOEε4* represents the major genetic risk factor for sporadic Alzheimer disease (AD), increasing disease risk in homozygous carriers by 12-fold as compared to carriers of the *APOEε3/ε3* genotype^16^.

So far, the physiological relevance of sortilin-mediated neuronal uptake of lipids in the human brain remains unclear. However, such functions may well provide a molecular explanation for the genome-wide association of *SORT1* with both AD^17^ and frontotemporal lobe dementia (FTD)^18^, the two most common forms of age-dependent neurodegeneration in the human population. Here, we used humanized mouse strains and iPSC-derived cell models to recapitulate the human brain lipid metabolism *in vivo* and *in vitro* and to elucidate the significance of sortilin-dependent lipid homeostasis for neuronal health. Our studies identified a unique metabolic concept whereby apoE-bound poly-unsaturated fatty acids (PUFA) internalized by sortilin are converted into ligands for peroxisome proliferator-activated receptor (PPAR) α, a key transcriptional regulator of enzymes involved in use of long-chain fatty acids as metabolic fuel for mitochondrial energy production ^19^. This sortilin pathway is essential to sustain neuronal energy homeostasis when use of glucose is limited, and it is lost in the presence of apoE4 that disrupts sortilin’s function as neuronal lipoprotein receptor.

## RESULTS

### Genetic or functional sortilin deficiency impacts brain fatty acid metabolism

To interrogate the significance of sortilin and apoE interaction for brain (lipid) metabolism, we initially tested the impact of *Sort1* and *APOE* genotypes on the brain metabolome in humanized mouse models. These mouse strains carried a targeted replacement of the murine *Apoe* locus with genes encoding human apoE3 (E3) or apoE4 (E4)^20^. In addition, the animals were either wildtype (WT) or genetically deficient (KO) for *Sort1*^1,14^. When exploring possible phenotypes related to the physiological interaction of sortilin with apoE, we focused on metabolic traits distinct in E3WT mice as compared to the other three genotypes (E3KO, E4WT, and E4KO). This strategy was based on our hypothesis that interaction of sortilin with apoE3 in E3WT animals sustains proper metabolism and neuroprotective actions of lipids (Fig. 1A). However, these lipid pathways will be lost in E3 mice genetically *null* for sortilin (E3KO). The same phenotypes, as in E3KO, should be seen in E4 mice, regardless of being WT or KO for *Sort1*, as these mouse models are either functional (E4WT) or genetically deficient (E4KO) for sortilin’s endocytic activity (Fig. 1A).

**Figure 1:**
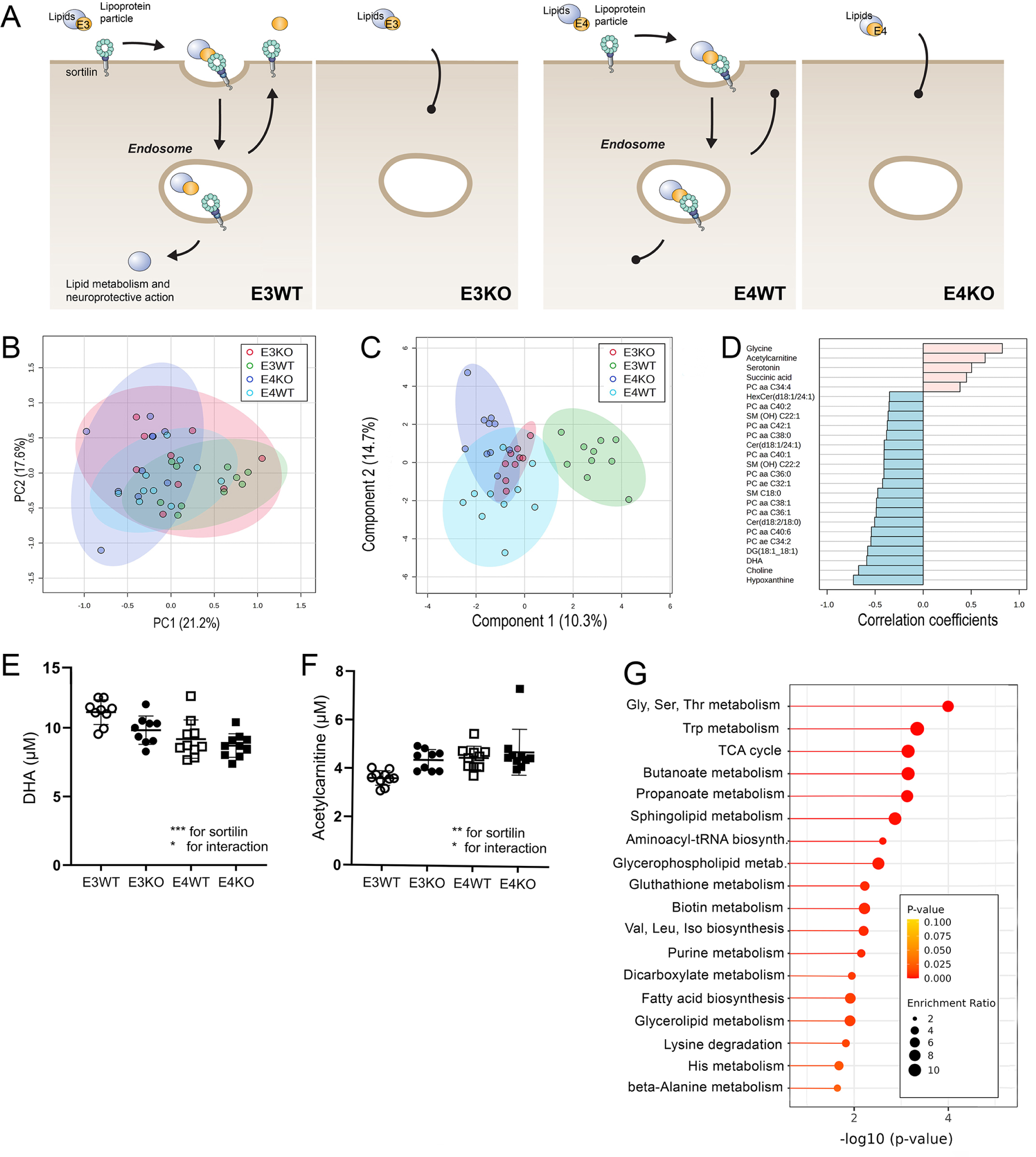
Global metabolomics identifies interaction of sortilin and apoE3 in short-chain fatty acid transport and metabolism by brain mitochondria. **(A)** Proposed model of sortilin and apoE isoform-specific interactions in neuronal lipid handling. Normal receptor function in cellular lipid homeostasis is seen in wildtype mice expressing apoE3 (E3WT), governing cellular uptake and neuroprotective metabolism and action of essential lipids. Functional (E4WT) or genetic (E3KO, E4KO) sortilin deficiency disrupts neuronal uptake and metabolism of such lipids in the other three genotypes. **(B, C**) Brain cortices from male mice of the indicated *Sort1* and human *APOE* genotypes (age 12 weeks, n=10 per genotype group) were subjected to targeted metabolomics using LC-MS and FIA-MS analysis (MxP Quant 500 kit, Biocrates Life Science). Data were analyzed using MetaboAnalyst 5.0 (https://www.metaboanalyst.ca/). Unsupervised Principal Component Analysis (B) and supervised Partial Least-Squares Discriminant Analysis (sparsed variant) (C) were used to interrogate separation of samples according to genotype (E3WT, green; E3KO, red; E4WT, light blue; E4KO, dark blue). The performance of the PLS-DA model was evaluated using 5-fold cross validation (classification error rate 23% for 2 components). (**D**) Pattern hunter analysis of brain metabolome data identifying metabolites characteristic of sortilin and apoE3 interaction with E3WT being different to the other three genotypes. The X-axis gives the calculated correlation coefficients, reflecting positive (pink) or negative (blue) correlation of metabolite concentration with the tested pattern. (**E - F**) Concentrations for DHA (E) and acetylcarnitine (F) in the data set are given (Two-way ANOVA; *, p<0.05; **, p<0.01; ***, p<0.001). DHA, docosahexaenoic acid. (**G**) Pathway analyses based on Metabolite Set Enrichment procedure identifying the most significantly altered metabolic pathways in the brain metabolome data set.

Accordingly, we used unbiased liquid chromatography mass spectrometry (LC-MS) and flow injection analysis (FIA)-MS based metabolomics to compare the concentrations of 630 polar and non-polar metabolites in brain cortices of our four mouse strains (MxP Quant 500 kit, Biocrates Life Science). Unsupervised Principal Component Analysis failed to document overt genotype-specific distinctions in global brain metabolism (Fig. 1B).

However, supervised partial least-squares discriminant analysis (sparse variant) identified differences in the E3WT brain metabolome compared to the other three genotypes as a major discriminator in the dataset (Fig. 1C). Pattern hunter analysis of the dataset listed a number of metabolites indicative of the distinction between E3WT and the other three genotypes, including docosahexaenoic acid (DHA), the most abundant and neuroprotective μ-3-PUFA in the brain. DHA levels were decreased in E3KO and in E4s as compared to E3WT (Fig. 1D and E). By contrast, levels of acetylcarnitine, involved in transport of C2 short-chain fatty acids (SCFA) were increased in E3KO and E4 brains as compared to E3WT (Fig. 1D and F). Metabolite Set Enrichment Analysis further supported changes in fatty acid metabolism as a major discriminator of E3WT versus the other three genotypes. Significantly discriminating biological terms included propanoate and butanoate metabolism, dicarboxylate metabolism, fatty acid biosynthesis, as well as tricarboxylic acid cycle, and metabolism of amino acids (Fig. 1G). Remarkably, no such link between *APOE* and *Sort1* genotypes was observed when performing unsupervised (Fig. S1A) or supervised (Fig. S1B) discrimination analyses of metabolome data obtained from plasma samples of the same mouse strains. Plasma metabolites discriminating E3WT from the other three genotypes did not include any fatty acids or related metabolites, but a number of triglycerides that are constituents of very low-density lipoproteins, but are of otherwise poorly characterized function (Fig. S1C). These changes were likely related to a role of sortilin in hepatic handling of lipoproteins^2,3,8,9^.

Taken together, unbiased metabolomics identified a specific interaction of sortilin and isoform apoE3 in control of fatty acid metabolism. This interaction was unique to the brain and not seen in plasma, despite prominent expression of sortilin and apoE in peripheral tissues, such as the liver.

### Loss of sortilin or the presence of apoE4 prevents use of long-chain fatty acids as metabolic fuel by neurons

Given the suggested impact of sortilin and apoE3 interaction on brain fatty acid transport and metabolism, we investigated genotype-specific consequences for energy homeostasis using real-time measurement of mitochondrial respiration (Seahorse Technology, Agilent). To specifically query neuronal respiration, we isolated synaptosomes from brain cortices of the mice. Synaptosomes are a source of neuronal mitochondria, entrapped during preparation from brain tissue by density gradient ultracentrifugation^21^. Comparing mitochondrial oxygen consumption rate (OCR), we identified a significantly higher maximal respiration in E3WT as compared to E3KO, E4WT, or E4KO synaptosomes, with no statistically significant differences between the latter three genotypes (Fig. 2A and D). By contrast, other mitochondrial parameters, including basal respiration (Fig. 2B), proton leak (Fig. 2C), or ATP-linked respiration (Fig. 2E) did not show this interaction of genotypes. Also, Western blot analysis of mitochondrial proteins in synaptosomal extracts as well as electron microscopy of thin sections of synaptosomes did not reveal any discernable differences in mitochondrial protein concentrations (Fig. S2A) or structural integrity of these organelles (Fig. S2B) between genotypes. In line with the functional interaction of sortilin and apoE3 being specific to the brain, real-time measurement of respiration in mitochondria isolated from the livers of these mice failed to show any genotype-dependent differences in mitochondrial activities (Fig. S2C-G).

**Figure 2:**
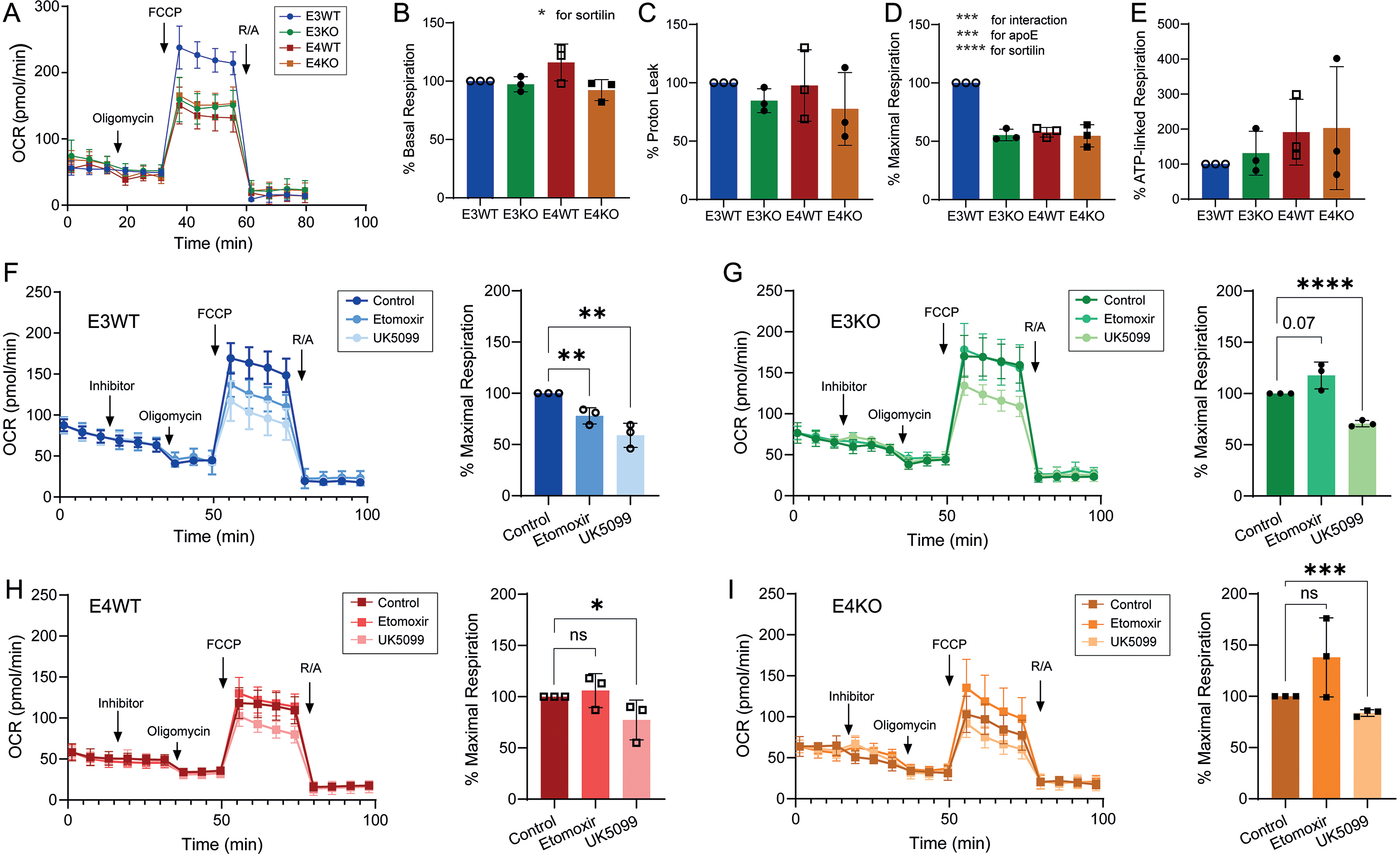
Loss of sortilin or the presence of apoE4 renders neuronal mitochondria insensitive to inhibition of long-chain fatty acid import. (**A**) Quantification of real-time cellular oxygen consumption rates (OCR) in synaptosomes isolated from brain cortices of male mice of the indicated *Sort1* and *APOE* genotypes (12 weeks of age). Representative respiration profiles of E3WT, E3KO, E4WT, and E4KO synaptosomes are given. OCR was measured under basal conditions (0 - 20 min) and following sequential addition of oligomycin, carbonyl cyanide-p trifluoromethoxyphenylhydrazone (FCCP), as well as rotenone and antimycin A (R/A) at the indicated time points. (**B - E**) Quantification of basal respiration (B), proton leak (C), maximal OCR (D), and ATP-linked respiration (E) in synaptosomes of the indicated genotypes relative to E3WT (set to 100%). Data are given as mean ± SD of n=3 biological replicates (mice) per genotype group. Each biological replicate data point is the mean of n=20-24 technical replicates. Statistical significance of data was tested using 2way ANOVA with Tukey’s multiple comparison test. (**F - I**) The dependency of mitochondrial respiration on availability of glucose or long-chain fatty acids (LCFA) was assessed as described under (A) in the presence of control buffer (control) or solutions containing 8 μM UK5099 or 16 μM etomoxir (inhibitor). For each genotype, one exemplary experiment is shown (to the left) as well as the quantification of % maximal OCR compared to the control condition (set to 100%; to the right). Data are given as mean ± SD from n=3 biological replicates. Each biological replicate data point is the mean of n=10-18 technical replicates. Statistical significance of data was tested using unpaired Student’s *t* test (two-tailed). *, p < 0.05; **, p < 0.01; ***, p < 0.001; ****, p < 0.0001).

Jointly, our data suggested a distinct impact of sortilin and apoE3 interaction on maximal respiratory capacity of neuronal mitochondria. To test whether a shortage of lipids as alternative metabolic fuel may limit the maximal activity of the respiratory chain in the E3KO and E4 synaptosomes, we performed a substrate oxidation stress test by treating synaptosomes with UK5099 or etomoxir. UK5099 targets the mitochondrial pyruvate carrier (MPC), blocking the ability of mitochondria to use glucose as metabolic fuel^22^. Etomoxir is an inhibitor of CPT1A, disrupting mitochondrial import of long-chain fatty acids (LCFA)^23^. Treatment of E3WT synaptosomes with UK5099 or etomoxir significantly reduced % maximal respiration as compared to buffer-treated samples (Fig. 2F), documenting the ability of neuronal mitochondria from E3WT to use both glucose and LCFA as energy substrates. By contrast, synaptosomal mitochondria from E3KO (Fig. 2G), E4WT (Fig. 2H), or E4KO (Fig. 2I) were sensitive to treatment with UK5099 but failed to respond to etomoxir. These findings argued for an inability of the latter three genotypes to utilize LCFA as alternative fuel for mitochondrial energy production.

To directly test the impact of genotypes on lipid fuel use in neurons, we performed real-time measurement of mitochondrial respiration in synaptosomes substituted with long-, medium-, or short-chain fatty acids. These studies were performed under conditions of limited supply with pyruvate to mimic a situation of brain stress imposed by glucose hypometabolism, as during aging or AD ^24–27^. Maximal respiration increased in E3WT synaptosomes when substituted with palmitate (Fig. 3A). However, no increase in maximal respiration was seen in E3KO (Fig. 3B), E4WT (Fig. 3C), or E4KO (Fig. 3D) synaptosomes upon palmitate supplementation. These data corroborated our assumption of an inability of the latter three genotypes to use LCFA as metabolic fuel. The situation was different when synaptosomes were provided with SCFA. Both addition of acetic acid or butyric acid significantly increased maximal respiration in E3KO (Fig. 3F), E4WT (Fig.3G), and E4KO (Fig. 3H), but no effect was seen in E3WT (Fig. 3E). We also tested the impact of heptanoic acid and octanoic acid, medium-chain fatty acids (MCFA) used in the treatment of patients with inherited deficiencies of import and oxidation of LCFA into mitochondria ^28^. Similar to the situation with SCFA, MCFA also increased maximal respiration, an effect seen in all genotypes (Fig. 3I-L). Jointly, these findings substantiated our model of a switch in metabolic fuel choice from LCFA in E3WT to SCFA and MCFA in E3KO, E4WT, and E4KO.

**Figure 3:**
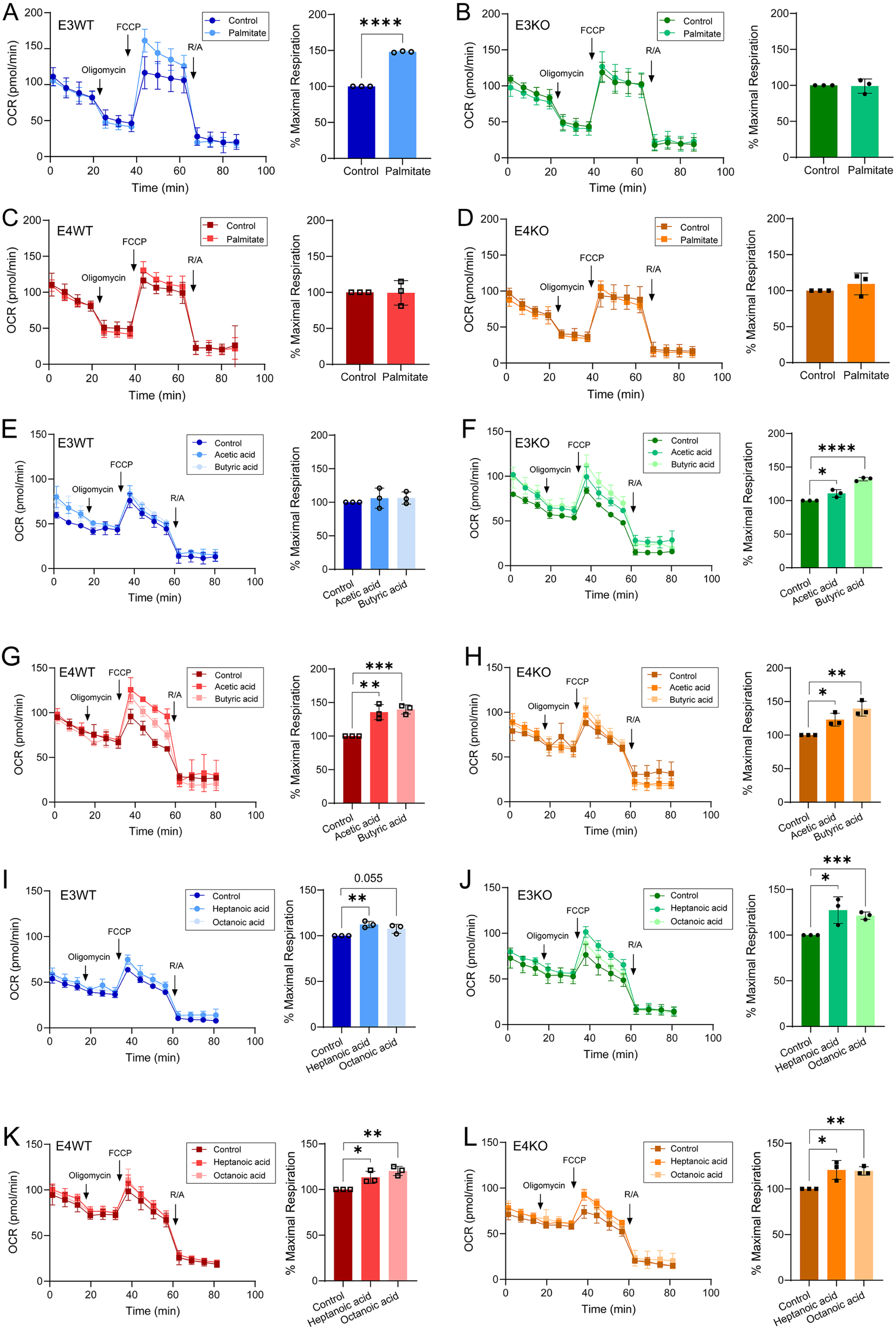
Loss of sortilin or the presence of apoE4 prevents consumption of long-chain fatty acids by neuronal mitochondria. Quantification of real-time cellular oxygen consumption rates (OCR) in synaptosomes isolated from brain cortices of male mice of the indicated *Sort1* and human *APOE* genotypes (12 weeks of age). OCR was measured under basal conditions (0 - 20 min) and following sequential addition of oligomycin, carbonyl cyanide-p-trifluoromethoxyphenylhydrazone (FCCP), as well as rotenone and antimycin A (R/A) at the indicated time points. (**A - D**) The dependency of mitochondrial respiration on long-chain fatty acids was assessed by pre-treatment of synaptosomes with 150 μM palmitate. (**E - H**) The dependency of mitochondrial respiration on short-chain fatty acids was assessed by pre-treatment of synaptosomes with 100 μM acetic acid or 100 μM butyric acid. (**I - L**) The dependency of mitochondrial respiration on medium-chain fatty acids was assessed by pre-treatment of synaptosomes with 200 μM heptanoic acid or 200 μM octanoic acid. For each genotype, one exemplary experiment is shown (to the left) as well as the quantification of % maximal OCR as compared to control treatment of the same genotype (set to 100%). Data are given as mean ± SD of n=3 biological replicates (animals). Each biological replicate is the mean of n=8-15 technical replicates. Statistical significance of data was tested using unpaired Student’s *t* test (two-tailed; *, p < 0.05; **, p < 0.01; ***, p < 0.001; ****, p < 0.0001).

### *SORT1* deficiency decreases the levels of poly-unsaturated fatty acids and endocannabinoids in human apoE3 neurons

To dissect the molecular mechanism underlying the inability of E3KO or E4 synaptosomes to use LCFA for neuronal respiration, we quantified global brain transcript levels of multiple genes implicated in fatty acid metabolism and mitochondrial lipid consumption. However, no genotype-dependent differences in expression levels were detected in any of the tested genes (Fig. S3). These findings argued that sortilin and apoE3 interaction distinctly impacts the cellular metabolism of neurons, not all brain cell types. This assumption was supported by the predominant expression of the receptor in neuronal cell types in the brain^29^.

To interrogate neuron-specific functions for sortilin in the human brain, we established iPSC-based cell models to recapitulate neuronal and glial interactions in human brain lipid homeostasis. To do so, we used endonuclease-based strategies to introduce *SORT1* gene defects into human iPSC lines homozygous for *APOEϕ.3* or *APOEϕ.4* (Fig. S4A-B). Loss of sortilin expression was confirmed by Western blotting (Fig. S4C). *APOE* and *SORT1* genotypes did not impact pluripotency of the iPSC lines as shown by testing expression of pluripotency markers using immunocytochemistry (Fig. S4E-F) and qRT-PCR (Fig. S4D, G-H).

To recapitulate cell type-specific interactions in human brain lipid homeostasis, we differentiated the four iPSC lines into astrocytes using an established protocol of retroviral-induced overexpression of *SOX9* and *NFIB* ^30^. The viral constructs also encoded mCherry to trace transduced cells (Fig. 4A). All four genotypes generated astrocytes (referred to herein as induced astrocytes, iAs) that were comparable in appearance as well as in expression of astrocyte markers vimentin and glial fibrillary acid protein (GFAP), documented by immunocytochemistry (Fig. 4B) and qRT-PCR (Fig. 4C). Minor differences were seen in transcript levels for S100β and aldehyde dehydrogenase family member L1 (ALDH1), but these changes did not show interaction between *APOE* and *SORT1* genotypes (Fig. 4C). Importantly, iAs of all four genotypes produced and secreted similar amounts of apoE variants as shown by comparative qRT-PCR (Fig. 4C) and Western blotting (Fig. 4D).

**Figure 4:**
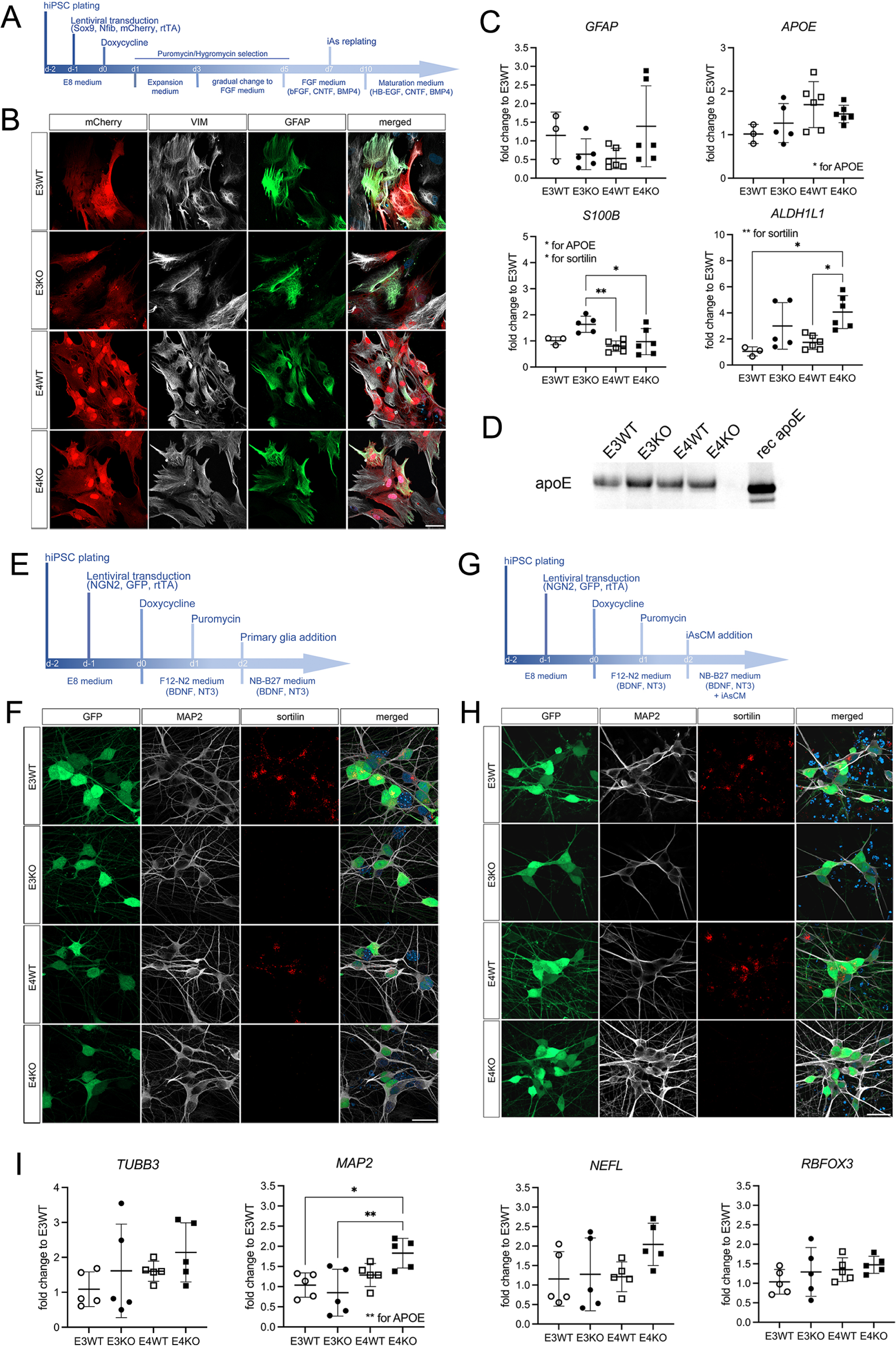
iPSC-based co-culture model of cell type-specific interactions in human brain lipid metabolism. **(A)** Protocol used for generation of human astrocytes (iAs) from induced pluripotent stem cells (hiPSCs). **(B)** Representative immunofluorescence images of human iAs of the indicated *SORT1* and *APOE* genotypes visualized for native mCherry (red) as well as immunostainings for vimentin (VIM, white) and glial fibrillary acidic protein (GFAP, green). Cells were counterstained with DAPI (blue, in the merged images). Scale bar: 50 μm. **(C)** Quantitative RT-PCR analysis of selected astrocyte marker genes in human iAs of the indicated *SORT1* and *APOE* genotypes (day 21 of culture). Individual biological replicates (n=3-6) from 2-3 individual differentiation experiments as well as mean ± SD of the entire genotype group are shown. Statistical significance of data was tested using 2-way ANOVA with Tukey’s multiple comparison test. *ALDH1L1*, aldehyde dehydrogenase family 1 member L1; *S100B*, S100 calcium binding protein beta. **(D)** Levels of secreted apoE isoforms were determined in 30 µl cell supernatant of human iAs (day 21 of culture) using Western blotting. Recombinant human apoE (rec apoE, 12.5 ng) served as detection control. **(E)** Protocol used for generation of hiPSC-derived human neurons (iNs) in co-culture with mouse primary glia. **(F)** Representative immunofluorescence images of human iNs of the indicated *SORT1* and *APOE* genotypes generated in co-culture with mouse primary glia (day 12 of culture). Neurons were visualized for native green fluorescent protein (GFP, green) and immunostained for microtubule associated protein 2 (MAP2, white) and sortilin (red). Cells were counterstained with DAPI (blue, in the merged images). Scale bar: 25 μm. **(G)** Protocol used to generate hiPSC-derived human neurons (iNs) by addition of iAs-conditioned medium (iAsCM). **(H)** Representative immunofluorescence images of human iNs of the indicated *SORT1* and *APOE* genotypes grown in the presence of iAsCM (day 12 of culture). Neurons were visualized for native GFP (green) and immunostained for MAP2 (white) and sortilin (red). Neurons were counterstained with DAPI (blue, in the merge images). Scale bar: 25 μm. **(I)** Quantitative RT-PCR analysis of selected neuronal marker genes in human iNs of the indicated *SORT1* and *APOE* genotypes (day 7 of culture). Individual biological replicates (n=5) of 3 individual differentiation experiments as well as mean ± SD of the entire genotype group are shown. Statistical significance of data was tested using 2-way ANOVA with Tukey’s multiple comparison test. *, p < 0.05; **, p < 0.01). *MAP2,* microtubule associated protein 2; *NEFL*, neurofilament light chain; *TUBB3*, class III beta tubulin; *RBFOX3*, RNA binding Fox-1 homolog 3.

In parallel, all four iPSC lines were differentiated into cortical neurons using virus-induced expression of *NGN2* ^31^. In addition, the retroviral constructs also encoded GFP as a transduction marker. Using established protocols described in the STAR method section, E3WT, E3KO, E4WT, and E4KO iPSC lines generated cortical neurons when co-cultured with primary mouse glia (Fig. 4E and F). To avoid the presence of glia in our neuronal cultures, we next adapted the neuronal differentiation protocol by adding media conditioned by human iAs instead of murine glia (Fig. 4G). This protocol reliably produced iPSC-derived neurons from all four genotypes (referred to as induced neurons, iNs), comparable in cellular appearance as well as in expression of neuronal markers TUJ1 (TUBB3), neurofilament light chain (NEFL), and RNA binding Fox-1 homolog 3 (RBFOX3) as shown by immunocytochemistry (Fig. 4H) and qRT-PCR (Fig. 4I). Minor differences in transcript levels were seen for MAP2, but these changes did not show an interaction between *APOE* and *SORT1* genotypes (Fig. 4I).

Initially, we used our iPSC-derived cell models to test whether interaction of sortilin with lipidated apoE governs lipid metabolism in human neurons, as shown in mouse models before ^14^. To do so, we cultured iNs of defined *SORT1* and *APOE* genotypes for 10 days in conditioned medium from iAs of the same genotypes and measured neuronal levels of PUFA, and their bioactive derivates endocannabinoids, by LC-MS/MS thereafter. Loss of sortilin in E3KO iNS resulted in a drastic decrease in cellular PUFA content when compared with the isogenic E3WT line (Fig. 5A). Among other lipids, levels of ω6- and ω3 PUFAs were reduced, as well as the levels of the endocannabinoid precursors linoleic acid (C18:2n-6) and DHA. Sortilin deficiency also decreased neuronal levels of endocanabinoids (eCB) and eCB-like metabolites, including 2-arachidonoylglycerol (2-AG), synaptamide, and linoleoyl ethanolamide (LEA), and caused a concomitant increase in levels of anandamide (Fig. 5A). Loss of PUFA and eCB content in E3KO neurons was not due to alterations in levels of lipids released by iAs as the concentrations of the tested lipids were comparable in conditioned media from E3WT and E3KO iAs (Fig. 5B). Importantly, *SORT1* deficiency did not alter the concentrations of most PUFA and eCBs in E4KO iNs when compared with isogenic E4WT neurons (Fig. S5A), a finding in agreement with apoE4 rendering WT neurons functionally deficient for sortilin.

**Figure 5:**
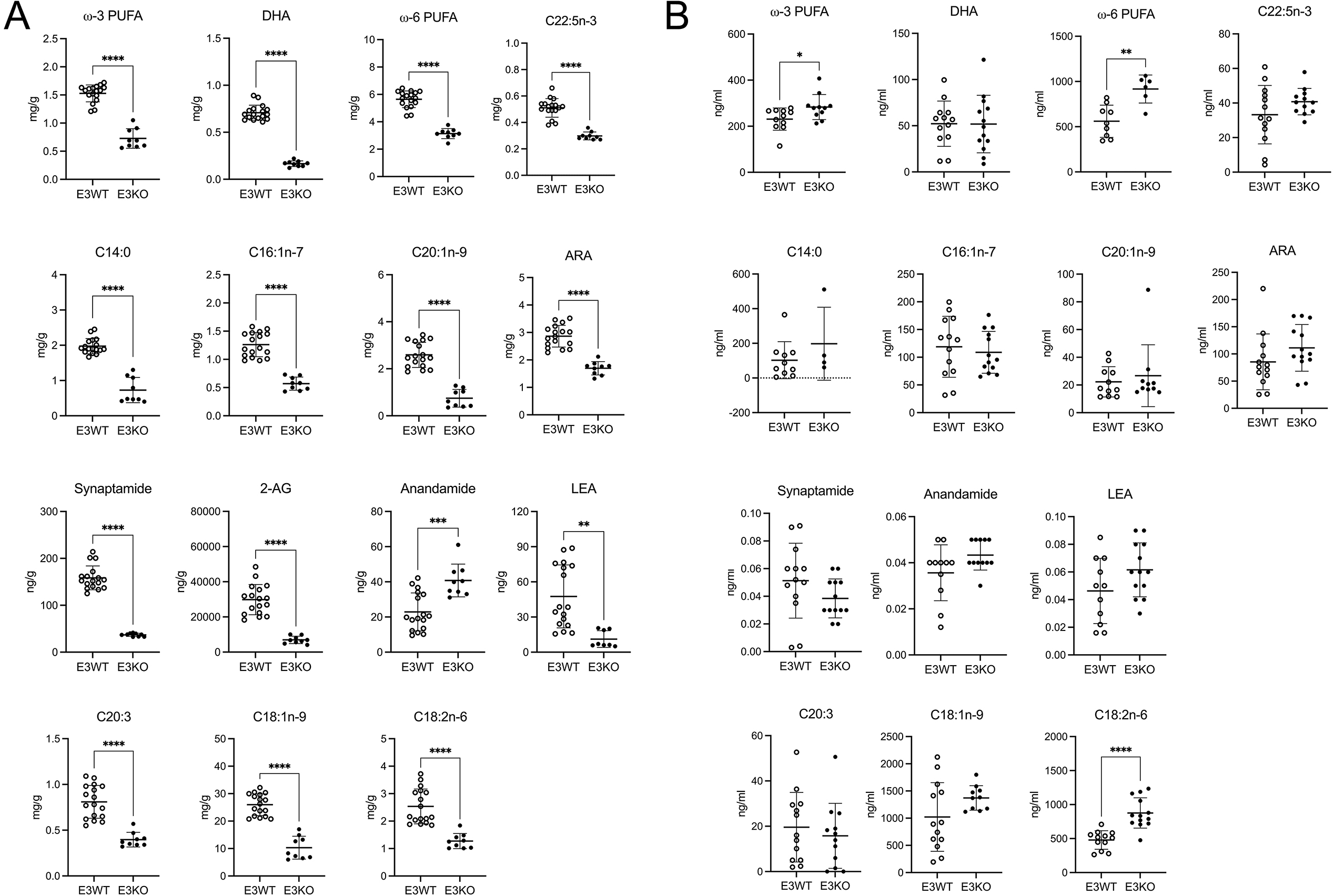
*SORT1* deficiency decreases the levels of poly-unsaturated fatty acids and endocannabinoids in human E3 neurons but not astrocytes. (**A**) Concentrations of selected lipids in iPSC-derived cortical neurons were determined using LC-MS. Neurons (day 12 of culture) were *APOEε3/ε3* and either wildtype (E3WT) or genetically deficient for *SORT1* (E3KO). Individual biological replicates (n=9-17) of 2 - 3 individual differentiation experiments are given. (**B**) Concentrations of selected lipids in the cell supernatant of iPSC-derived E3WT and E3KO astrocytes (days 21 - 28 of culture) were determined using LC-MS. Individual biological replicates (n=13) of 4 - 6 individual differentiation experiments are shown. Individual data points as well as mean ± SD of the entire genotype group are shown. Statistical significance of data was tested using unpaired Student’s *t* test (two-tailed; *, p < 0.05; **, p < 0.01; ***, p < 0.001; ****, p < 0.0001). ARA, arachidonic acid; 2-AG, 2-arachidonoylglycerol; DHA, docosahexaenoic acid; LEA, linoleoyl ethanolamide; PUFA, poly-unsaturated fatty acid.

### Impaired LCFA fuel use in sortilin-deficient neurons is rescued by induction of neuronal PPARα activity

eCBs and eCB-like metabolites exert their neuroprotective functions through various molecular mechanisms, including by acting as ligands for transcription factors of the peroxisome proliferator-activated receptor (PPAR) gene family ^32,33^. To query the consequence of decreased PUFA and eCB levels in E3KO iNs, we performed comparative expression profiling of PPAR target genes in E3WT and E3KO iNs. Using a microarray-based strategy to assess transcript levels of 84 PPAR targets, we identified 30 transcripts that were dysregulated in E3KO when compared to isogenic E3WT iNs (Fig. 6A-B). Most of these genes showed reduced transcript levels when neurons lacked sortilin (Fig. 6C). With relevance to lipid homeostasis, decreased transcript levels included those encoding long-chain acyl-CoA dehydrogenase (*ACADL*), involved in fatty acid β-oxidation, carnitine transporter SLC22A5, as well as fatty acid transport proteins SLC27A1, 5, and 6. Decreased transcription was substantiated by qRT-PCR for selected PPAR targets, such as fatty acid translocase (*CD36*) and *ACADL* (Fig. 6D). Sortilin deficiency in E3KO iNs also decreased transcript levels of *PPARA* (encoding PPARα), a master transcriptional regulator of cellular lipid metabolism (Fig. 6D) ^19^. No impact on transcription was seen for genes encoding PPAR family members PPAR8 and PPARψ (Fig. 6D). Importantly, no effect of sortilin deficiency on any of the above genes was seen comparing transcription profiles in isogenic E4KO and E4WT iNs, once more corroborating functional sortilin deficiency in the presence of apoE4 (Fig. S6).

**Figure 6:**
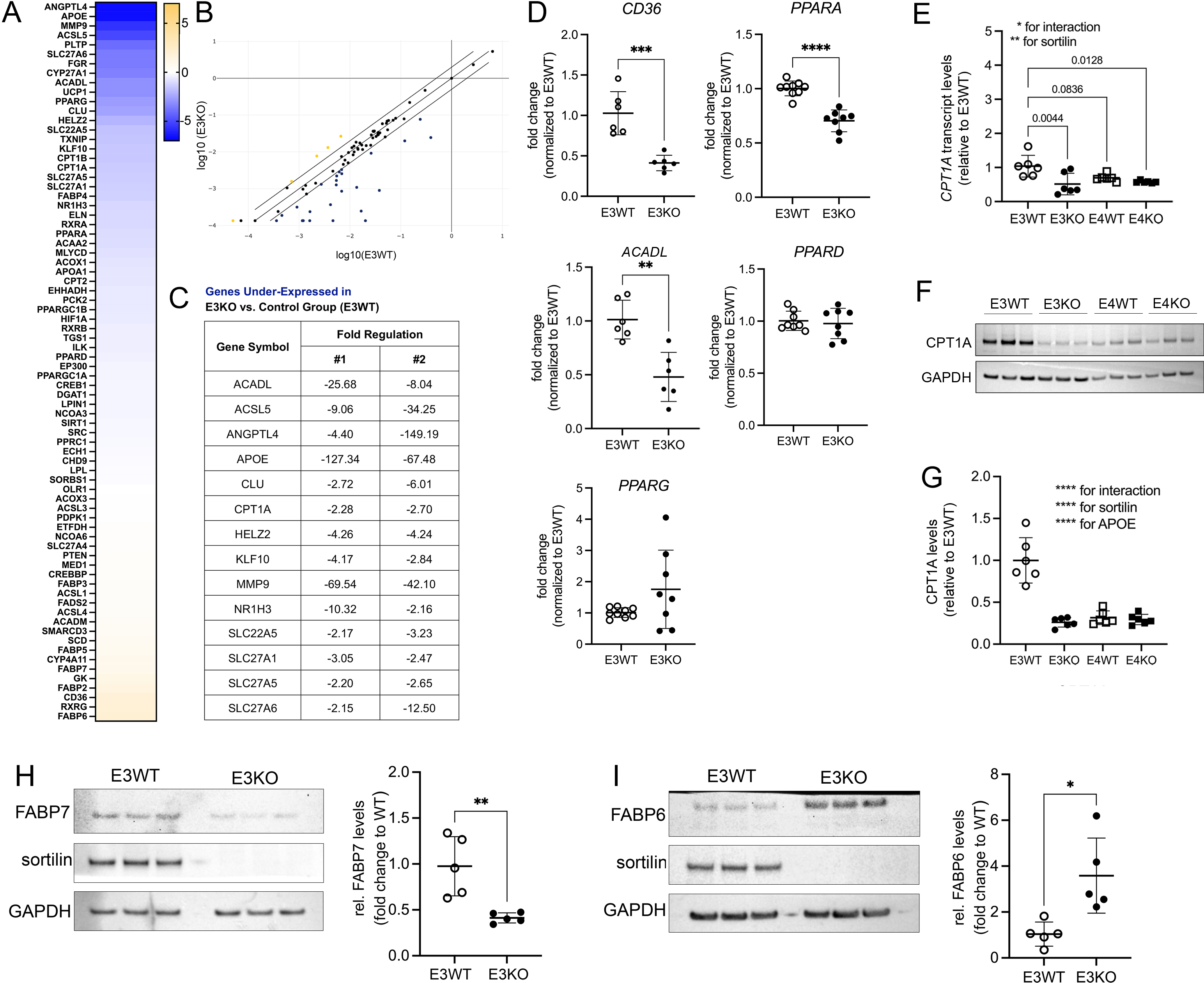
Sortilin deficiency disrupts PPAR-dependent gene transcription and expression of fatty acid binding protein 7 in apoE3 iNs. **(A-C)** Expression levels of 84 peroxisome proliferator-activated receptor (PPAR) target genes in iPSC-derived neurons treated with iPSC-derived astrocyte conditioned media were tested using the Human PPAR Targets RT^2^ Profiler PCR Array. Neurons (day 12 of culture) were *APOEε3/ε3* and either wildtype (E3WT) or genetically deficient for *SORT1* (E3KO). One exemplary experiment from a total of n=2 individual biological replicates is shown. (A) Expression levels are given as heat map with levels in E3KO presented as log_2_ fold change compared to E3WT (set to 0). The blue and yellow color spectra indicate down- and up-regulated genes, respectively. (B) Scatterplot analysis comparing log-transformed relative expression for all tested PPAR target genes in E3WT and E3KO neurons. The center diagonal line indicates unchanged gene expression, while the outer diagonal lines indicate 2-fold regulation threshold. Genes with >2-fold difference in transcript level between groups are presented as blue and yellow dots, indicating down- or up-regulated genes, respectively. (C) Selected list of genes downregulated by >2 fold in E3KO compared to E3WT neurons. Data from two independent experiments are shown. **(D)** Quantitative RT-PCR analysis of the indicated PPAR target genes in E3WT and E3KO human neurons treated with iAs-conditioned medium (day 12 of culture). Individual biological replicates (n=6-9) of 2 - 3 individual differentiation experiments are shown. Statistical significance of data was tested using unpaired Student’s *t* test (two-tailed). **(E)** *CPT1A* transcript levels in human neurons of the indicated *SORT1* and *APOE* genotypes treated with iAs-conditioned medium were tested using qRT-PCR. Individual data points from n=6 biological replicates (2 - 3 individual differentiations) as well as mean ± SD of the entire genotype group are shown. Statistical significance of data was tested using 2-way ANOVA with Tukey’s multiple comparison test. (**F - G)** Levels of CPT1A protein were determined in total lysates of human neurons (12 day of culture) of the indicated *SORT1* and *APOE* genotypes treated with iAs-conditioned medium using Western blotting (F) and densitometric scanning of replicate blots thereof (G). Detection of GAPDH served as loading control. Data in (G) represent data points from individual biological replicates (n=6) of 2 - 3 individual differentiation experiments. Individual data points as well as mean ± SD of the entire genotype group are shown and given as relative to E3WT (set to 1). Statistical significance of data was tested using 2-way ANOVA. **(H - I)** Levels of fatty acid binding protein (FABP) 7 (H) and FABP6 (I) as determined by Western blotting in total lysates from human neurons of the indicated *SORT1* and *APOE* genotypes. For each panel, one exemplary Western blot as well as quantitative analysis from densitometric scanning of replicate blots are shown. Values represent the mean ± SD of n=5 biological replicates (from 2 individual differentiations). Statistical significance of data was tested using unpaired Student’s *t* test (two-tailed). *, p < 0.05; **, p < 0.01; ***, p < 0.001; ****, p < 0.0001.

One transcriptional target of PPARα with particular relevance to mitochondrial lipid handling is CPT1A, the mitochondrial importer for LCFA and target of etomoxir. Levels of *CPT1A* transcript (Fig. 6E) and CPT1A protein (Fig. 6F-G) were significantly higher in E3WT as compared to E3KO, E4WT, or E4KO iNs, with no statistically significant difference between the later three genotypes. Loss of PPARα-induced expression of CPT1A provided a possible molecular explanation for the insensitivity of E3KO or E4 synaptosomes to etomoxir. Alterations in fatty acid handling in neurons lacking sortilin were also substantiated when testing for expression of fatty acid binding proteins (FABP) in E3WT and E3KO cells. Loss of sortilin coincided with reduced levels of FABP7, a known sortilin interactor^15^ and intracellular carrier for PUFA and eCB (Fig. 6H), and with a concomitant increase in levels of FABP6, an alternative intracellular fatty acid carrier^34^ (Fig. 6I).

To elucidate the molecular mode of sortilin and apoE3 interaction in energy homeostasis in human neurons, we compared mitochondrial respiration in E3WT, E3KO, E4WT, and E4KO iNs supplemented with conditioned medium from iAs of the corresponding genotypes. Maximal respiration was always compared in neuronal cultures untreated (control) or treated with UK5099 or etomoxir. Mirroring phenotypes seen in murine synaptosomes, maximal respiration decreased in E3WT iNs when treated with UK5099 or etomoxir (Fig. 7A). By contrast, maximal respiration in E3KO iNs (Fig. 7B), in E4WT iNs (Fig. 7C), or in E4KO iNs (Fig. 7D) was reduced by UK5099, but not by etomoxir. These findings confirmed the inability of the later three genotypes to use LCFA as alternative metabolic fuel in human neurons as well. As with murine synaptosomes, these metabolic alterations were not due to changes in mitochondrial volume (Fig. S7A and B), content of key mitochondrial proteins (Fig. S7C), or structural architecture of mitochondria (Fig. S7D) in human neurons.

**Figure 7:**
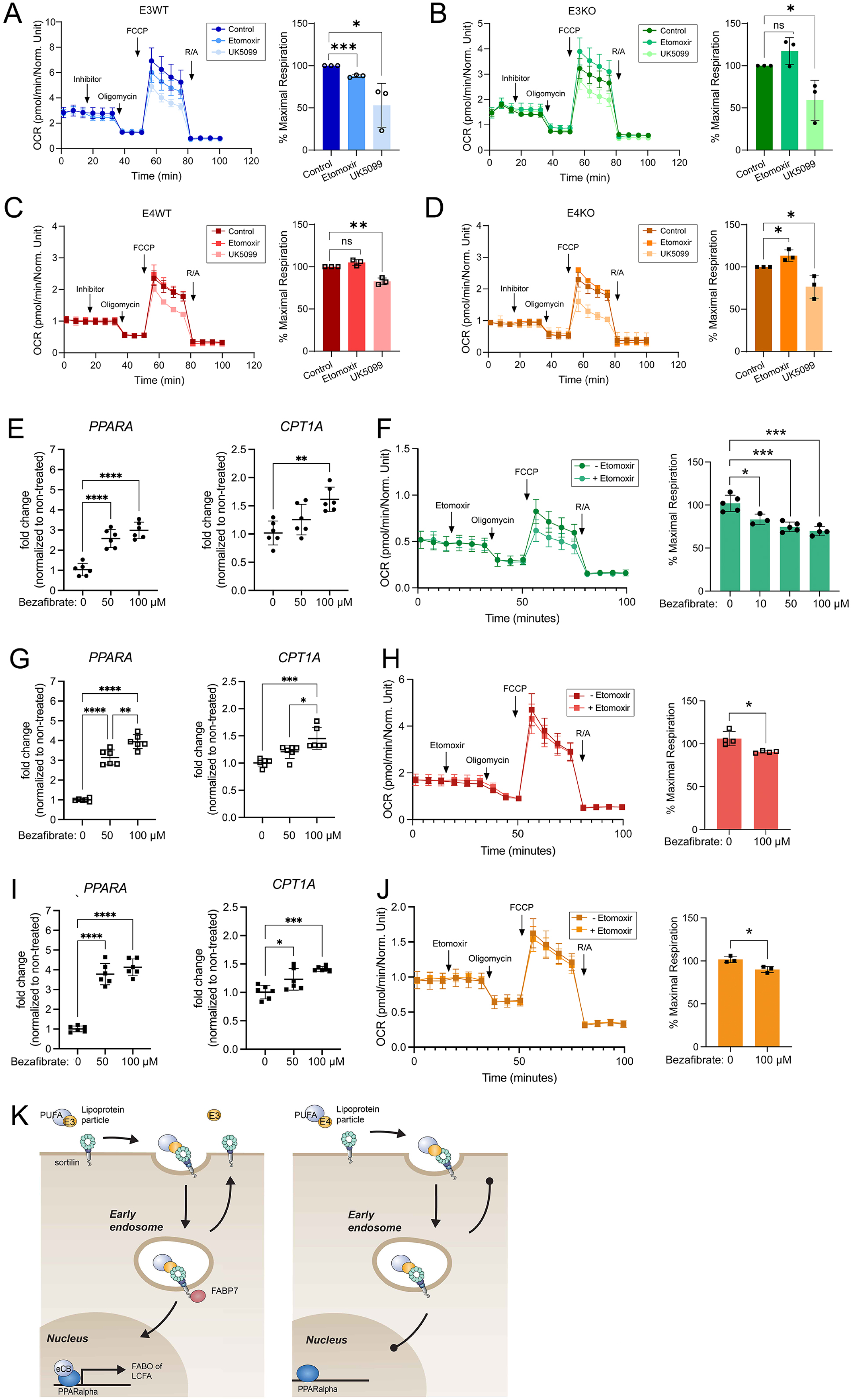
Rescue of use of long-chain fatty acids as metabolic fuel in human E3KO and E4 neurons by PPARα agonist bezafibrate. (**A - D**) Quantification of real-time cellular oxygen consumption rates (OCR) in human neurons E3WT (A), E3KO (B), E4WT (C), or E4KO (D). OCR was measured under basal conditions (0 - 20 min) and following sequential addition of oligomycin, carbonyl cyanide-p trifluoromethoxyphenylhydrazone (FCCP), as well as rotenone and antimycin A (R/A). In addition, neurons had been treated with control buffer or solutions containing 8 μM UK5099 or 16 μM etomoxir at the indicated time points (inhibitor). For each genotype, one exemplary experiment is shown as well as the quantification of % maximal OCR as compared to the control buffer treatment of the same genotype for the entire data set (set to 100%). Data represent the mean ± SD of n=3 biological replicates. Each biological replicate is the mean of n=10-15 technical replicates. Statistical significance of data was tested using unpaired Student’s *t* test (two-tailed). (**E**) Transcript levels of *CPT1A* and *PPARA* in human E3KO neurons were tested using qRT-PCR. Cells have not been treated (0) or pretreated with the indicated concentrations of bezafibrate. Individual data points from n=6 biological replicates (3 individual differentiations) as well as mean ± SD of the entire genotype group are shown. Statistical significance of data was tested using One-way ANOVA with Bonferroni’s multiple comparisons test. (**F**) The dependency of mitochondrial respiration on PPARα activity was assessed by pre-treatment of human E3KO iNs with bezafibrate for 48 hours before substrate oxidation stress test in the presence of control buffer or a solution containing 16 μM etomoxir (as detailed in panels A-D above). One exemplary experiment using 100 µM bezafibrate in the presence or absence of etomoxir is shown as well as the quantification of % maximal respiration as mean ± SD from n=3 biological replicates (n=10-15 technical replicates) of etomoxir-treated versus non-etomoxir treated E3KO neurons in the absence or presence of the indicated concentrations of bezafibrate. Statistical significance of data was tested using unpaired Student’s t test (two-tailed). (**G - J)** The dependency of mitochondrial respiration on PPARα activity was assessed by pre-treatment of human E4WT (G and H) or E4KO (I and J) neurons with bezafibrate for 48 hours. Panels G and I document *CPT1A* and *PPARA* transcript levels as tested in (E) above. Panels H and J depict substrate oxidation stress tests in the presence of control buffer or a solution containing 16 μM etomoxir (as detailed in panels A-D above). One exemplary experiment using 100 µM bezafibrate in the presence or absence of etomoxir is given as well as the quantification of % maximal respiration as mean ± SD from n=3 biological replicates (n=8-12 technical replicates) of etomoxir-versus non-etomoxir treated E4 neurons in the absence or presence of bezafibrate. Statistical significance of data was tested using unpaired Student’s t test (two-tailed). **(K)** Model of sortilin and apoE isoform-specific interaction in facilitation of PPARα-dependent mitochondrial respiration of long-chain fatty acids. This pathway is operable with apoE3 (left schematic) but lost with apoE4 (right schematic). E, apoE; FABO, fatty acid beta oxidation; FABP7, fatty acid binding protein 7; eCB, endocannabinoids, LCFA, long-chain fatty acids; PUFA, poly-unsaturated fatty acids. *, p < 0.05; **, p < 0.01; ***, p < 0.001; ****, p < 0.0001.

To query whether loss of lipid-induced transcriptional activity of PPARα may be responsible for the inability to metabolize LCFA, we treated E3KO iNs with the PPARα agonist bezafibrate ^35^. As expected, induction of PPARα activity by bezafibrate increased transcript levels of its targets *PPARA* and *CPT1A* in E3KO iNS in a dose-dependent manner (Fig. 7E). Importantly, bezafibrate treatment also conferred sensitivity to etomoxir in E3KO iNS as documented by a bezafibrate dose-dependent decrease in maximal respiration in cells treated with etomoxir as compared to control iNs (Fig. 7F). This phenotype contrasted the inability of E3KO iNS to respond to etomoxir in the absence of bezafibrate (see Fig. 7B). Similar to the situation in E3KO, bezafibrate treatment also increased PPARα activity in E4WT and E4KO iNs (Fig. 7G and I) and it conferred sensitivity to etomoxir (Fig. 7H and J).

In conclusion, our data identified a novel concept in neuronal energy homeostasis and the central role of the lipoprotein receptor sortilin in this process (Fig. 7K). In detail, we propose that endocytic uptake of lipidated apoE3 by this receptor delivers PUFA to neurons to be converted into ligands for the transcriptional activator PPARα. As shown in established cell lines before, apoE3 uptake also requires the interaction of sortilin with the carrier FABP7, facilitating intraneuronal lipid transport^15^. PPARα-dependent induction of metabolic gene expression profiles enables neurons to import LCFA into mitochondria and use it as alternative metabolic. However, this pathway is lost in the presence of apoE4 that disrupts sortilin’s endocytic activity, resulting in impaired PPARα activation. As a consequence, neurons exposed to apoE4 are unable to efficiently utilize LCFA as metabolic fuel. Instead, they revert to PPARα-independent mitochondrial import of SCFA and MCFA, energetically less favorable substrates to support mitochondrial energy production.

## DISCUSSION

Our data suggest a novel concept in control of metabolic fuel choice in neurons that depends on cellular uptake of PUFA transported by apoE. This lipid uptake pathway requires the action of sortilin, a lipoprotein receptor genetically associated at a genome-wide level with neurodegenerative disorders AD and FTD. Conceptually, sortilin’s role in governing metabolic adaptations in neurons may be particularly relevant in conditions when the availability of glucose becomes limited.

### Sortilin, a facilitator of alternative metabolic fuel choice in neurons

Healthy neurons rely on glucose for energy production, contributing to a significant portion of the metabolite consumed by the human body. However, availability of glucose becomes limited during brain aging ^24,26,36–39^. Among other mechanisms, a marked decrease in expression of glucose transporters contributes to glucose hypometabolism in neurons and other brain cell types ^36,39–43^. Reduced metabolism of glucose is also a feature of the brain in AD ^44–46^ and FTD ^47,48^, aggravating the deterioration of neuronal functions. Impaired availability of glucose in the aging brain enforces a switch in neurons to alternative metabolic fuels, including ketone bodies ^49^ and fatty acids ^50^. The ability of neurons to use LCFA as metabolic fuel is documented by findings in this study that blockade of mitochondrial uptake by etomoxir reduces maximal respiration in E3WT synaptosomes and iNS by 15-20%. Based on the loss of sensitivity to etomoxir, this alternative metabolic fuel choice is absent in E3KO and E4 mitochondria and coincides with a reduction in maximal respiration as compared to E3WT (Fig. 2D). Finally, when pyruvate is limited, E3WT synaptosomes respond to exogenous fatty acid supplementation with a substantial increase in maximal respiration. This effect is seen with palmitate, but not with acetic or butyric acid, indicating that neuronal mitochondria express an enzyme repertoire that favors mitochondrial import of LCFA over SCFA (Fig. 3). Although our analyses rely on measurement of neuronal respiration *in vitro*, unbiased metabolomics of humanized mouse models support our hypothesis of the sortilin/apoE3 axis to regulate use of fatty acids by mitochondria. This conclusion is evidenced by induced expression of acetylcarnitine (Fig. 1F) and other adaptations in fatty acid metabolism (Fig. 1G) in mice lacking sortilin activity (i.e., E3KO, E4WT, and E4KO).

### Sortilin promotes the neuronal activity PPARα, a regulator of fatty acid metabolism

What may be the molecular pathway whereby sortilin impacts alternative metabolic fuel choices in neurons? Using targeted lipidomics, we show that loss of sortilin activity in human E3KO neurons coincides with a drastic reduction in neuronal levels of PUFA and their bioactive derivates, most notably endocannabinoids and endocannabinoid-like metabolites (Fig. 5A). These findings recapitulate data shown by us in mouse models before^14^. This PUFA defect is attributed to a prominent role of the receptor in neuronal uptake of lipidated apoE, providing essential lipids for cellular lipid metabolism and action^10,14^. Several apoE receptors have been documented in neurons, including the low-density lipoprotein (LDL) receptor, the LDL-receptor related protein 1, and the apoE receptor 2 (reviewed in ^12^). Loss of sortilin activity in gene-targeted mice increases circulating levels of apoE in the brain, due to impaired cellular clearance of the carrier^10^. Sortilin deficiency also reduces apoE uptake in primary mouse neurons by approximately 70%^10^. Such pronounced clearance defects have not been reported in other apoE receptor KO models. As well as by quantity, sortilin may also distinguish itself from other apoE receptors by functionality as it interacts with FABP7^15^. FABP7 is a brain-specific lipid chaperone for cytosolic trafficking of PUFA and eCBs ^51–53^. FABP7-dependent trafficking is required for biosynthesis of eCBs from PUFA, and for their delivery to nuclear PPARs for regulation of gene transcription ^54–57^. Sortilin is essential for proper sorting and stable expression of FABP7 in established cell lines^15^, a mechanism now corroborated for human neurons in this study (Fig. 6H). Thus, sortilin is in a unique position amongst various apoE receptors to link extracellular (i.e., apoE) and intracellular (i.e., FABP7) PUFA transport processes in neurons.

Given the drastic decrease in PUFA and eCB levels in E3KO iNs as compared to isogenic E3WT cells, impaired activation of PPARα is the likely reason for alterations in mitochondrial lipid handling in this genotype. This conclusion is supported by decreased levels of synaptamide, 2-AG, and LEA (Fig. 5A), known ligands of this transcription factor ^58,59^. Impaired activity of PPARα in human neurons lacking sortilin is substantiated by the reduced expression of multiple transcriptional targets with relevance to lipid handling, most notably CPT1A, the mitochondrial importer for LCFA (Fig. 6A-G). Importantly, the insensitivity to etomoxir seen in E3KO iNs can be rescued by bezafibrate, proving impaired PPARα activity as the underlying molecular cause of this defect (Fig. 7F).

The metabolism of neurons is intimately linked to that of astrocytes which support neurons with apoE-bound lipids and other metabolites. Recent studies have documented dysregulation in astrocyte metabolism in response to aging or apoE4 ^60,61^. While our studies do not exclude possible consequences of sortilin deficiency or the presence of apoE4 on astrocyte metabolism, alterations in neuronal PUFA and eCB homeostasis shown here are not due to reduced secretion of these lipids or their precursors from iAs (Figs. 5B and S5B). The hypothesis of such defects being inherent to receptor dysfunction in neurons is supported by the lack of sortilin and apoE3 interactions in the plasma metabolome (Fig. S1).

PUFA and eCB have numerous functions in cell metabolism and action. Thus, reduced fatty acid levels in neurons lacking sortilin possibly impact multiple cellular activities related to lipid homeostasis, or even unrelated receptor actions, such as in neurotrophin signaling^1,62,63^. This prediction is in line with changed levels of polar metabolites distinguishing E3WT from E3KO and E4 brain metabolomes, such as glycine, serotonin, and choline (Fig. 1D). Such changes may represent secondary consequences of neuronal dyslipidemia or reflect the loss of receptor activities, other than lipoprotein uptake, not investigated as yet.

### ApoE4 renders neurons functionally deficient for sortilin

*SORT1* shows genome-wide association both with AD^17^ and FTD^18^, the two most common forms of age-related dementias in patients. This association argues for a crucial function of the receptor in healthy aging of the brain. Sortilin’s role in FTD is linked to its ability to act as clearance receptor for progranulin, a major etiologic agent in FTD^64–66^, while its mode of action in AD remains unclear. As both dementias are characterized by a hypometabolism of glucose ^44–48,67^, deteriorating lipoprotein receptor functions of sortilin in the aging brain may well aggravate deficits in neuronal energy production in the elderly.

An additional mechanism specific to AD concerns the detrimental impact of apoE4 on sortilin activity. Similar to other receptors bound by apoE ^68,69^, binding of apoE4 traps internalized sortilin in early endosome, preventing discharge of cargo and recycling to the cell surface^15^. This noxious propensity of apoE4 is due to an increase in isoelectric point, as compared to apoE3, that results in unfolding of the protein in the acidic milieu of endosomes ^70–74^. In our studies, phenotypes in E4WT were indistinguishable from that of E3KO, including alterations in the brain metabolome, in mitochondrial malfunction, and in PPARα deficiency. These findings strongly argue for loss of sortilin activity in the brain of *APOEϕ4* carriers, an assumption supported by identical features seen in E4 mouse brains and human neurons, regardless of being WT or KO for the receptor gene. Multiple pathological mechanisms have been reported for apoE4 (reviewed in ^75–77^), and functional inactivation of sortilin adds another mechanistic explanation to the detrimental actions of the most important genetic risk factor for sporadic AD known to date.

In conclusion, using mouse and iPSC-derived cell models of the human brain lipid metabolism, we have deciphered a central role for the lipoprotein receptor sortilin in control of neuronal energy metabolism, a function likely to contribute to the impact of this receptor on aging brain health and disease seen by human genetics.

## Acknowledgement

We are indebted to T. Pasternack, K. Kampf, H. Zweers, C. Schiel, K.-M. Pedersen, and A. Højland for expert technical assistance. Studies were funded in part by the European Research Council (BeyOND no. 335692), the Alzheimer Forschung Initiative (#18003), and the Novo Nordisk Foundation (NNF18OC0033928) to TEW.

## Author contributions

AKG, JPG, EZP, RFG, IMR, and NST designed and conducted the experiments and analyzed data. CG provided essential expert advice. JK, SK, MR, SD, PB, and TEW conceptualized the study and evaluated data. AKG, JPG, PB, and TEW wrote the manuscript.

## MATERIALS AND METHODS

### Mouse models

The generation of mouse strains carrying a targeted replacement of the murine *Apoe* locus with human *APOEe3* or *APOEe4* ^20^, and being wildtype (*Sort1^+/+^*) or genetically deficient for *Sort1* (*Sort1^-/-^*)^78^ has been described^14^. The animals were kept on a normal chow (4.5% crude fat, 39% carbohydrates). All animal experimentation was conducted in male mice on an inbred C57Bl6/J background following approval by local ethics committees (X9017/17).

Animals were 12 weeks of age.

### Generation and culture of gene targeted induced pluripotent stem cell lines

Human induced human pluripotent stem cell (iPSC) lines either *APOEε3/ε3* (https://hpscreg.eu/cell-line/BIHi005-A) or *APOEε4/ε4* (https://hpscreg.eu/cell-line/WTSIi009-A) were obtained from the MDC Technology Platform Pluripotent Stem Cells (in-house) or the Welcome Trust Sanger Institute, respectively, and designated as E3WT and E4WT lines in this study. Stem cells were cultured under feeder-free conditions on Matrigel-coated plates (Corning) using Essential 8 Flex medium (Gibco). Cells were passaged using 0.5 mM EDTA solution when the culture reached 80% confluency. To obtained single cell suspensions, the cells were treated with StemPro Accutase (Gibco) and replated in medium supplemented with 10 µM of Rock inhibitor Y27632 (Cayman Chemical) for the first 24 hours. Cell lines were regularly tested negative for mycoplasma.

To generate sortilin-deficient isogenic subclones of BIHi005-A (designated E3KO), the *SORT1* gene was targeted using transcription activator-like effector nucleases (TALENs) essentially as described^79^. In brief, TALEN constructs were designed to target the *SORT1* start codon using TALEN Targeter webtool (Cornell University, https://tale-nt.cac.cornell.edu/node/add/talen). Four TALENs segments were digested from an 832-plasmid library (Addgene) using BsmBI and ligated into a TALENs vector carrying a FokI endonuclease domain, and either GFP or RFP (*SORT1* ATG left-GFP: I: CGGC, II: GGCA, III: TTCG, IV: GCG; *SORT1* ATG right-RFP: I: CCGC, II: AGCT, III: CCCC IV: AGG). TALEN pairs were transfected into BIHi005-A iPSC cells using Lipofectamine 3000 (Thermo Fisher) according to manufacturer’s instructions. Cells were subjected to FACS sorting 48 hours post-transfection based on GFP and RFP expression. The double-positive cells were replated at low density in Essential 8 medium to form single cell colonies. Colonies were expanded and genomic DNA extracted for PCR amplification the ATG region of *SORT1* (forward primer: 5’-CGTTCCAGCCAATCAGTCCC-3’; reverse primer: 5’- AGCTTGGCGACGAAGTCC-3’). PCR products were analyzed on 2% agarose gels to identify insertions or deletions, followed by Sanger sequencing to verify a homozygous disruption of the ATG. Positive clones were quality controlled for karyotype stability and cryopreserved.

To generate sortilin-deficient isogenic subclones of HPSI0913i-diku_1 cells (named E4KO), we utilized the CRISPR/Cas9 genome-editing system as described in STAR protocols^80^. In brief, a single guide RNA (sgRNA) targeting exon 4 of *SORT1* was designed using the Synthego knockout guide design tool (https://design.synthego.com). The sgRNA sequence (5’-GCCAAATTCAGTCCGAATAA-3’) obtained from Integrated DNA Technologies (https://eu.idtdna.com) and complexed with Hi.Fi Cas9 to form ribonucleoprotein (RNP) particles, which were transfected into HPSI0913i-diku_1 cells using the Neon transfection 10 µl kit (Thermo Fisher Scientific), according to the manufacturer’s instructions. Editing efficiency was estimated 48 hours post-transfection by Sanger sequencing, and data were analyzed using the Synthego ICE tool. To isolate single-cell clones, we used the automated Iota Sciences IsoCell platform. Targeted clones were identified by Sanger sequencing and inference of CRISPR edit analysis, quality controlled for karyotype stability, and tested regularly negative for Mycoplasma.

### Global metabolomics

Metabolomics of murine brain cortex and plasma samples was performed using the MxP Quant 500 kit (Biocrates Life Science AG). This kit is an assay based on phenylisothiocyanate (PITC) derivatization of the target analytes using internal standards for quantitation. It enables quantitative analysis of up to 630 metabolites from 26 analyte classes, including small molecules (1 alkaloid, 1 amine oxide, 20 amino acids, 30 amino acid related, 14 bile acids, 9 biogenic amines, 1 carbohydrate, 7 carboxylic acids, 1 cresol, 12 fatty acids, 4 hormones, 4 indoles and derivatives, 2 nucleobases and related and 1 vitamin) and lipids (40 acylcarnitines, 76 phosphatidylcholines, 14 lysophosphatidylcholines, 15 sphingomyelins, 28 ceramides, 8 dihydroceramides, 19 hexosylceramides, 9 dihexosylceramides, 6 trihexosylceramides, 22 cholesteryl esters, 44 diglycerides and 242 triglycerides; full list at https://biocrates.com/wp-content/uploads/2022/02/biocrates-Quant500-list-of-metabolites-v6-2022.pdf). All reagents, internal and calibration standards, quality controls, and filter plates were provided by the manufacturer.

Male mice of the indicated *Sort1* and human *APOE* genotypes were used for the experiment (n=10 animals per genotype). Mice were fasted for 6 hours before EDTA-blood was collected from the submandibular vein, centrifuged to obtain plasma, and stored at - 80°C. Brain cortices were dissected and also stored at -80°C. Before analysis, tissues were weighed, transferred into vials containing ceramic beads (Precellys), and mixed with cold MeOH as extraction solvent at a ratio of 6:1 (µl/mg tissue). Samples were lysed using a Precellys 24 tissue homogenizer (Precellys) combined with a nitrogen cooling unit (Cryolys, Precellys). Assay plate preparation was performed according to the manufacturer’s protocols. Briefly, 10 µl tissue lysate or 20 µl plasma were transferred to the upper microtiter plate and dried under a nitrogen stream. Thereafter, 50 µl of a 5% PITC solution was added. After incubation, the filter spots were dried before the metabolites were extracted using 5 mM ammonium acetate in MeOH (300 µl) into the lower 96-well plate after further dilution using running solvent A. Measurements were based on liquid chromatography mass spectrometry (LC-MS) or flow injection analysis (FIA)-MS techniques for polar metabolites and lipids, respectively. Quantification was carried out using internal standards (for LC and FIA) and a calibration curve for specific metabolites (LC). The LC-MS system was comprised of a 1290 Infinity ultrahigh performance liquid chromatography system (Agilent, Santa Clara, CA, USA), coupled to a QTrap 5500 (AB Sciex GmbH, Darmstadt, Germany) with a TurboV source. Evaluation of the instrument’s performance prior to sample analysis was carried out by system stability tests using provided test mixtures for LC-MS and FIA-MS. Gradient conditions are indicated in Tabs. 1 and 2 for LC and FIA, respectively. Acquisition method parameters are shown in Tab. 3. All compounds were identified and quantified using isotopically-labeled internal standards and multiple reaction monitoring for LC and full MS for FIA. Optimized and raw data were computed in Met*IDQ*^TM^ version Oxygen (Biocrates Life Science AG).

An in-house developed script was used for data quality analysis and pre-processing (Metaquac)^81^. Filtering was done based on Biocrates internal QC2 with MetIDQ status “valid” and minimum 60% valid values per group. Compounds with RSD >15% in QC2 were removed from the dataset. For samples, also MetIDQ status “valid” with minimum 60% valid values (6/9) was used. In total (mean of each biological group), data on 157 compounds from 39 cortex samples and 406 compounds from 36 plasma samples were exported into the MetaboAnalyst 5.0 platform (https://www.metaboanalyst.ca/). Features with more than 40% missing values were removed and remaining missing values were replaced by the k-nearest neighbors method. Samples were normalized by median and data were log transformed and pareto-scaled. One-way ANOVA with false discovery rate correction for multiple comparisons was performed for pre-selecting features that differed between genotype groups. To explore the data, unsupervised principal component analysis (PCA), supervised partial least squares discriminant analysis (sPLS-DA), as well as Pattern Hunter tool were applied. PCA was performed in MetaboAnalyst using the prcomp package with calculations based on singular value decomposition. sPLS-DA is a clustering technique that maximizes the covariance between groups. It uses multivariate regression techniques to extract information predicting class membership through linear combination of original variables. MetaboAnalyst performs the sPLS-DA with the algorithm, provided by R package. Performance of obtained model was evaluated using 5-fold cross validation, and classification error was calculated.

Metabolite Set Enrichment Analysis (MSEA) is a way to identify biologically meaningful patterns that are significantly enriched in quantitative metabolomic data. A compound concentration table was used as input data and Over Representation Analysis (ORA) was implemented using the hypergeometric test to evaluate whether a particular metabolite set was represented within the given compound list more than expected by chance. The enrichment ratio was derived by dividing the number of hits in a specific metabolic pathway divided by the expected number of hits. One-tailed *p* values were provided after adjusting for multiple testing. The analysis was based on metabolite set KEGG human metabolic pathways library (https://www.genome.jp/kegg/pathway.html).

### Bioenergetic assays in murine liver mitochondria

Mice were sacrificed by cervical dislocation and liver dissected on ice. Liver tissues were homogenized in isolation buffer (225 mM mannitol, 75 mM sucrose, 0.1 mM EGTA, 10 mM HEPES-KOH (pH 7.4)) including 0.5% w/v fatty acid free bovine serum albumin using a drill-driven tissue homogenizer (IKA, EUROSTAR 20 digital) with 10 strokes at 200 rpm. The homogenate was centrifuged at 500g for 5 min at 4℃ and mitochondria were precipitated from the supernatant by additional centrifugation at 14,000g for 10 min at 4℃. The pellet was resuspended in 12% Percoll in isolation buffer. The supernatant was directly applied onto a 21% Percoll and centrifuged at 18,000g for 15 min at 4℃. Isolated mitochondria were obtained as a pellet and washed twice in isolation buffer, followed by centrifugation at 18,000g for 5 min at 4℃. The pellet was resuspended in assay buffer (220 mM mannitol, 70 mM sucrose, 10 mM EGTA, 10 mM KH_2_PO_4_, 5 mM MgCl_2_*7 H_2_O, 10 mM HEPES-KOH (pH 7.4)). Mitochondrial protein concentration was determined by Pierce Bradford Protein Assay Kit (Thermo Fisher Scientific).

Ten µg of mitochondrial protein in a volume of 30 µl were added to each well of a Seahorse XFe96/XF Pro Cell Culture Microplates (Agilent Technologies) coated with polyethylenimine (Sigma-Aldrich, P3143). The mitochondria loaded XF microplate was centrifuged at 1,900g for 20 min at 4℃ for adherence of the synaptosomes. Assay media (10 mM HEPES-KOH, 220 mM mannitol, 70 mM sucrose, 10 mM EGTA, 10 mM KH_2_PO4, 5 mM MgCl_2_*7H_2_O) with 0.2% FA-free BSA, 10 mM succinate and 2 mM malate were added to a final volume of 180 µl. Once the probe calibration was completed, the probe plate was replaced by the XF microplate. Two baseline measurements of oxygen consumption rates (OCR) were performed prior to sequential injection of 4 μM ADP, 2.5 μM oligomycin (Sigma-Aldrich, 75351), 4 μM FCCP (Sigma-Aldrich, C2920), and 4 μM antimycin A (Sigma-Aldrich, A8674) in combination with 4 μM rotenone (Sigma-Aldrich, 557368). OCR were measured by Seahorse XFe96 Flux Analyzer using a mix (30 s) and measure (3 min) cycle. Each sample was measured in at least 20 technical replicates. All experiments were conducted independently at least three times. Basal respiration was calculated as the differences between the last OCR measurement before oligomycin injection and the minimum OCR measurement after addition of rotenone and antimycin A. Maximal respiration was calculated as the differences between maximum OCR measurement after FCCP injection and the minimum OCR measurement after the addition of rotenone and antimycin A. ATP-linked respiration was determined as the differences by the last OCR measurement before oligomycin injection and the minimum OCR measurement after oligomycin injection. Proton leak was determined as the difference between minimum OCR measurement after oligomycin injection and the minimum OCR measurement after the addition of rotenone and antimycin A.

### Bioenergetic assays in synaptosomes

Mice were sacrificed by cervical dislocation and the brain cortices dissected on ice. The cortices were manually homogenized in gradient buffer (0.32 M sucrose, 5 mM Tris-HCl, 5 mM EDTA, 0.25 mM DTT) using a glass Dounce homogenizer (Bellco Glass, 1984-10002). Nuclear material was removed by centrifugation at 1,000g for 10 min at 4℃. The supernatant was applied onto a Percoll density gradient with 23%, 10% and 3% Percoll in gradient buffer. The gradient was centrifuged at 29,000g for 15 min at 4℃ using a Sorvall SS-34 rotor. The gradient fraction at the interface between the 23% and 10% Percoll containing synaptosomes were collected and protein concentration determined by Pierce Bradford Protein Assay Kit (Thermo Fisher Scientific, 23200).

Twenty µg of synaptosomal protein in a volume of 80 µl were added to each well of a Seahorse XFe96/XF Pro Cell Culture Microplates (Agilent Technologies, 103792-100) coated with polyethylenimine (Sigma-Aldrich, P3143). The mitochondria loaded XF microplate was centrifuged at 1,900g for 20 min at 4℃ for adherence of the synaptosomes. Pre-warmed assay media (2 mM MgSO_4_*7 H_2_O, 1.2 mM Na_2_SO_4_, 1.3 mM CaCl_2_, 0.4 mM KH_2_PO_4_, 3.5 mM KCl, 12 mM NaCl) containing 0.4% BSA, 10 mM sodium pyruvate, 10 mM glucose, and 2 mM glutamine was added to a final volume of 180 µl. Once the probe calibration was completed, the probe plate was replaced by the XF microplate. Three baseline measurements of oxygen consumption rate (OCR) were measured prior to sequential injection of 6 μM oligomycin, 4 μM FCCP, and 2 μM antimycin A in combination with 2 μM rotenone (Sigma-Aldrich, 557368). OCR were measured by Seahorse XFe96 Flux Analyzer using a mix (30 s), wait (2 min), and measure (3 min) cycle. Each sample was measured at least 20 technical replicates. All experiments were conducted independently at least three times. The respiratory parameters were calculated as described for liver mitochondria above.

### Substrate oxidation stress test in synaptosomes

To dissect endogenous substrate consumption, 20 µg of synaptosomal proteins were added to each well in a volume of 80 µl of a Seahorse XFe96/XF Pro Cell Culture Microplates (Agilent Technologies, 103792-100) coated with polyethylenimine (Sigma-Aldrich, P3143). The mitochondria-loaded XF microplate was centrifuged at 1,900g for 20 min at 4℃ for adherence of the synaptosomes. Pre-warmed assay media (2 mM MgSO_4_*7 H_2_O, 1.2 mM Na_2_SO_4_, 1.3 mM CaCl_2_, 0.4 mM KH_2_PO_4_, 3.5 mM KCl, 12 mM NaCl) containing 0.4% BSA, 10 mM sodium pyruvate, 10 mM glucose, and 2 mM glutamine was added to a final volume of 180 µl. Once the probe calibration was completed, the probe plate was replaced by the XF microplate. Three baseline measurements of oxygen consumption rate (OCR) were measured prior to injection of 8 μM UK5099 (Sigma-Aldrich, PZ0160), 16 μM etomoxir (Sigma-Aldrich, E1905), or assay media as control. Thereafter, sequential injection of 6 μM oligomycin, 4 μM FCCP, and 2 μM antimycin A in combination with 2 μM rotenone followed. OCR were measured by Seahorse XFe96 Flux Analyzer using a mix (30 s), wait (2 min), and measure (3 min) cycle. OCR for each genotype were measured independently three times. Each sample was measured in at least 10 technical replicates. Within each genotype group, OCR was normalized to the first measurement in the assay and maximal respiration reported as a comparison to the control. Maximal respiration was calculated as the differences between maximum OCR measurement after FCCP injection and the minimum OCR measurement after the addition of rotenone and antimycin A.

For measurement of exogenous fatty acid oxidation, 20 µg of synaptosomal proteins were added to each well of a Seahorse XFe96/XF Pro Cell Culture Microplates coated with Polyethylenimine. The samples were transferred in substrate-limited assay media (2 mM MgSO_4_*7 H_2_O, 1.2 mM Na_2_SO_4_, 1.3 mM CaCl_2_, 0.4 mM KH_2_PO_4_, 3.5 mM KCl, 12 mM NaCl) containing 0.4% BSA, 1 mM sodium pyruvate, 2 mM glucose, 1 mM glutamine, and 1 mM carnitine (Sigma-Aldrich, C0283). The mitochondria-loaded XF microplate was centrifuged at 1,900g for 20 min at 4℃ for adherence of the synaptosomes, followed by starvation for 30 minutes at 37℃. For assessment of long-chain fatty acid consumption, palmitate-BSA substrate (Cayman Chemical, 29558) or BSA (Cayman Chemical, 29556) was added to a final concentration of 300 μM palmitate. For assessment of medium-chain fatty acid consumption, heptanoic acid (Sigma-Aldrich, 75190) or octanoic acid (Sigma-Aldrich, C2875) were added at a final concentration of 200 μM. For assessment of short-chain fatty acid consumption, acetic acid (Sigma-Aldrich, 338826) or butyric acid (Sigma-Aldrich, B103500) were added to the final concentration of 100 μM. Assay media alone was used as control for all experiments. Once the probe calibration was completed, the probe plate was replaced by the XF microplate. Three baseline measurements of oxygen consumption rate (OCR) were measured prior to sequential injection of 6 μM oligomycin, 4 μM FCCP, and 2 μM antimycin A in combination with 2 μM rotenone. OCR were measured by Seahorse XFe96 Flux Analyzer using a mix (30 s), wait (2 min), and measure (3 min) cycle. All experiments were conducted independently at least three times with each sample measured in at least 8 replicates. The maximal respiration was reported as a comparison to the BSA or assay media treatment within each genotype group. Maximal respiration was calculated as the differences between maximum OCR measurement after FCCP injection and the minimum OCR measurement after addition of rotenone and antimycin A.

### Substrate oxidation stress test in iPSC-derived human neurons

On day 7 of differentiation, induced cortical neurons (iNs) were treated with accutase and plated as dissociated cell suspension in a density 1 x 10^5^ cells/well in Matrigel-coated Seahorse XFe96/XF Pro 96-well microplates using 100 ul/well of Seahorse XF DMEM medium supplemented with 10 mM glucose, 10 mM sodium pyruvate, 2 mM glutamine, and 10 µM Rock inhibitor. Once the probe calibration was completed, the probe plate was replaced by the XF microplate. Three baseline measurements of oxygen consumption rate (OCR) were carried out prior to injection of 8 μM UK5099 or 16 μM etomoxir. Seahorse XF DEMEM assay medium was used as control. Thereafter, 6 μM oligomycin, 4 μM FCCP, and 2 μM antimycin A in combination with 2 μM rotenone were injected sequentially. OCR were measured by Seahorse XFe96 Flux Analyzer using a mix (30 s), wait (2 min), and measure (3 min) cycle. OCR for each genotype were measured independently at least three times with each sample being measured in at least 8 technical replicates. The total cell numbers in the XF microplate wells were determined by fluorescence imaging with Hoechst on a designated area of the well using BioTek Instrument Cytation 5. Within each genotype group, respiration was normalized to the first measurement and the maximal respiration was reported as a comparison to the control condition. Maximal respiration was calculated as the differences between maximum OCR measurement after FCCP injection and the minimum OCR measurement after the addition of rotenone and antimycin A.

To test the impact of the PPARα agonist bezafibrate (Sigma-Aldrich, B7273) on neuronal respiration, the drug was dissolved in dimethyl sulphoxide (DMSO) to 10 mM stock solution that was stored at -20°C. On day 5 of neuronal differentiation, half of the cell medium was replaced with medium containing bezafibrate to achieve a final concentration of 10, 50, or 100 µM. After 48 hours of incubation, iNs were dissociated with accutase and plated in Matrigel-coated Seahorse XFe96/XF Pro (Agilent) 96-well microplates at a density 1.5 x 10^5^ iNs/well, and subjected to substrate oxidation stress test using 16 μM etomoxir as described above.

### Statistical analysis of bioenergetic assay data

Seahorse XFe96 Flux Analyzer data were processed in Wave 2.6.1 software (Agilent Technologies) following the manufacturer’s guidelines. Raw data were exported from the Wave Software to Prism 10 (GraphPad) for statistical analysis. The number of samples per group is referring to independent analyses (individual animals or neuronal cell cultures) indicated in the corresponding figure legends. The unpaired, two-tailed Student’s *t* test was used to determine statistical significance between the mean values of two groups. Two-way ANOVA with Tukey’s post-hoc test was used to compare the mean values of all four genotypes.

### Electron microscopy

Neurons and isolated synaptosomes were fixed at room temperature for 2 hours with 4% (w/v) formaldehyde, 2.5% (v/v) glutaraldehyde (Sigma-Aldrich G5882) in 0.1 M HEPES pH 7.4, followed by 2 days of fixation at 4°C. Thereafter, the samples were washed with Milli-Q water and pellets, obtained by centrifugation for 5 min at 300g, were embedded in 3% (w/v) aqueous agarose (Sigma-Aldrich A4018). Agarose-stabilized samples were cut into 0.5-1 mm^3^ cubes and prepared for electron microscopy.

Neuronal samples were processed using a modified version of the rOTO protocol as described (https://www.protocols.io/view/preparation-of-biological-tissues-for-serial-block-36wgq7je5vk5/v2). In brief, the samples were postfixed on ice for 1 hour with osmium tetroxide reduced with potassium ferrocyanide in 0.1M sodium cacodylate buffer pH 7. The final concentration was 1% (v/v) osmium tetroxide and 1.5% (w/v) potassium ferrocyanide in 0.1M sodium cacodylate buffer. The reduced osmium solution was washed with Milli-Q water and the samples incubated in 0.1% (w/v) aqueous thiocarbohydrazide. After rinsing with water, the samples were incubated for 30 min at room temperature in 1% (v/v) osmium tetroxide. Final contrast was achieved by incubating in 2% (w/v) uranyl acetate for 30 min at room temperature. After dehydration through a graded series of acetone, embedding was done in Durcupan ACM resin (Sigma-Aldrich). The blocks were polymerized for 48 hours at 60°C. Synaptosome samples were postfixed for 2 hours at room temperature using 1% (v/v) aqueous osmium tetroxide. Then, the samples were rinsed with Milli-Q water and incubated for 2 hours at 4°C in 2% (w/v) uranyl acetate. Dehydration using a graded ethanol series followed by propylene oxide incubation and resin infiltration with Polybed812 epoxy resin (Polysciences). Blocks were polymerized for 48 hours at 60°C.

Ultrathin 80 nm resin sections were sectioned using a Reichert Ultracut S ultramicrotome and picked up on Formvar-coated transmission electron microscopy grids. Reynolds Lead Citrate 3% (Delta Microscopies) was used for post-staining of the sections. Synaptosomes were imaged using a Zeiss EM910 80 kV Transmission Electron Microscope equipped with a 11M Quemesa CCD camera (EMSIS). Neurons were imaged using the Helios 5 Hydra CX Dual Beam system (Thermo Scientific, The Netherlands) operating at 2kV and 0.4 nA beam current. Micrographs were recorded using the CBS detector for back-scattered electrons and acquisition was done with the MAPS software package, version 3.27 (Thermo Scientific, The Netherlands).

### Generation of human iPSC-derived astrocytes (iAs)

Human astrocytes were generated from iPSCs as described ^30^. Briefly, iPSCs were dissociated with accutase and replated at a density of 5 x 10^5^ cells/well in Matrigel-coated 6-well plates using E8 Flex media supplemented with 10 µM Rock inhibitor (Y27632; Cayman Chemical). One day later (day -1), lentivirus vectors were added in fresh E8 Flex medium containing 8 μg/ml Polybrene (Millipore). The next day (day 0), the medium was replaced with E8 Flex medium containing doxycycline (2.5 μg/ml; Cayman Chemical), which was kept in the medium until the end of the experiment. On days 1 and 2, cells were cultured in expansion medium (DMEM/F12, 1% N2, 1% Glutamax, and 10% FBS, containing 1.25 μg/ml puromycin and 200 μg/ml hygromycin to select for cells containing lentivirus vectors expressing *Sox9* and *NfiB* (provided by the MDC Technology Platform Pluripotent Stem Cells). Days 3 - 5, the expansion medium was gradually exchanged to FGF medium (Neurobasal medium containing 2% B27, 1% NEAA, 1% Glutamax, 1% FBS, and 8 ng/ml human FGF (Peprotech), 5 ng/ml human CNTF (Peprotech) and 10 ng/ml human BMP4 (Peprotech). On day 6, the cell medium was replaced with fresh FGF medium before the cells were dissociated with accutase and replated in 2 x 10^5^ cells/well on Matrigel-coated 6-well plates on day 7 using FGF medium (supplemented with 10 µM Rock inhibitor from day 8). From days 10 to 17, half of the medium was exchanged every second day with maturation medium (1:1 DMEM/F12 and Neurobasal medium containing, 1% N2, 1% Glutamax, 1% sodium pyruvate, 5 μg/ml N-acetyl cysteine, 500 μg/ml dbcAMP, 5 ng/ml EGF-like growth factor, 10 ng/ml CNTF, and 10 ng/ml BMP4 (Peprotech)). On days 18 and 19, the medium was switched to NB-B27 (Neurobasal medium containing 2% B27 and 1% Glutamax). Astrocytes were kept in culture for up to 30 days, by replacing half of the medium every 2 days.

Conditioned media to supplement iPSC-derived human neurons was generated from astrocytes between day 21 and 28 of culture by collecting half of cell medium from each well every second day. Media samples from multiple wells collected the same day were pooled and centrifuged at 300g for 5 min to remove cell debris. Before adding to neurons, the conditioned medium was supplemented with 10 ng/ml human BDNF and 10 ng/ml human NT3, and diluted 1:1 with NB-B27 medium. If not used immediately, conditioned media were stored at 4°C for a maximum time of 3 days prior to addition to induced neurons.

### Generation of human iPSC-derived neurons

Induced neurons (iNs) were generated from iPSCs as described previously ^31^. Briefly, iPSCs were dissociated with accutase and replated at a density of 5.5 x 10^4^ cells/cm^2^ in Matrigel-coated 24- or 12-well plates or 4.3 x 10^4^ cells/cm^2^ in Matrigel-coated 6-well plates and 100 mm-dishes using E8 Flex media supplemented with 10 µM Rock inhibitor (Y27632; Cayman Chemical). Next day (day -1), iPSCs were transduced with lentivirus vectors in fresh E8 Flex medium containing 7 μg/ml Polybrene (Millipore). One day after infection (day 0), the medium was replaced with F12-N2 (DMEM/F12, 1% N2, 1% NEAA (Gibco)), containing 2 μg/ml doxycycline, 10 ng/ml human BDNF (R&D Systems), 10 ng/ml human NT3, and 0.2 μg/ml mouse laminin (Thermo Fisher). Doxycycline was retained in the medium through the experiment. On day 1, the medium was replaced with fresh F12-N2 medium containing 1 μg/ml puromycin to select vector expressing *Ngn2* (kindly provided by T. Sudhof, Standford Medicine). At day 2, astrocyte-conditioned medium (iAsCM), prepared as described above, was added and the medium exchanged with fresh iAsCM every second day. Alternatively, on day 2, primary glia dissociated in NB-B27 medium were added and half of the medium replaced every 2-3 days. iNs were kept for up to 12 days in culture.

### Lipid analyses of cultured human neurons

For fatty acid profiling, 30 mg of cell lysates were hydrolyzed with 100 µl 10 mol/l NaOH within 60 min at 80°C in presence of BHT. The samples were neutralized with 100 µl acetic acid (58 %). A 50 µl aliquot was diluted 1:10 with methanol containing deuterated internal standards (C12:0-d23, C18:0-d35, C18:2-d4, C20:4-d11, C20:5-d5, C22:0-d43, C22:6-d5, C24:0-d4; 10 ng each; Cayman Chemical, Ann Arbor MI). HPLC measurements were performed using an Agilent 1290 HPLC system with binary pump, autosampler and column thermostat equipped with a Phenomenex Kinetex-C18 column 2.6 µm, 2.1 x 150 mm column (Phenomenex, Aschaffenburg) using a solvent system of acetic acid (0.05 %) and acetonitrile. The solvent gradient started at 5 % acetonitrile and was increased to 98 % within 23 min and kept until 26 min with a flow rate of 0.4 ml/min and 1 µl injection volume. The HPLC was coupled with an Agilent 6470 triplequad mass spectrometer with electrospray ionisation source and operated in negative MRM mode. At least two mass transitions were detected for each fatty acid.

For eCB profiling, 10 mg cell lysates were homogenized in 300 µl distilled water. Then, 25 µl citric acid (0.4 mol/l), 10 µl of internal standards (10 µg/ml DHA-Ethanolamid-D4, Anandamid-D4, EPA-Ethanolamid-D4; Cayman Chemical, Ann Arbor MI) and 1 ml ethyl acetate were added. The samples were shaken for 10 min, centrifuged for 3 min at 3500 rpm and the upper phase recovered. Next, 1 ml ethyl acetate were added to the lower (sample) phase, followed by shaking for further 10 min and centrifugation for 3 min at 3500 rpm. Again, the upper phase was recovered and both supernatants combined. The solvent was removed to dryness at 40°C under a stream of N_2_, before the residue were resuspended in 100 µl acetonitrile and eCB measured. HPLC-measurement was performed using an Agilent 1290 HPLC system with binary pump, autosampler, and column thermostat equipped with a Agilent Poroshell EC120-C18 column 2.7 µm, 2.1 x 100 mm column (Agilent, Santa Clara, CA) using a solvent system of ammonium format (5 mM) and formic acid (0.05 %) in water and methanol. The solvent gradient started at 5 % methanol and was increased to 95 % within 11 min under gradient conditions and kept until 15 min with a flow rate of 0.4 ml/min and a 5 µl injection volume. The HPLC was coupled with an Agilent 6495 triplequad mass spectrometer with ESI source operated in positive MRM mode. Two mass transitions per compound were detected. All concentrations were calculated using Agilent Mass Hunter Software with individual calibration curves for each compound. All standards were purchase from Cayman chemical (Ann Arbor, MI).

### Protein expression analysis

Mouse synaptosomes or iPSC-derived cell types were lysed in RIPA buffer (50 mM Tris-HCl pH 7.4, 150 mM NaCl, 0.5% sodium deoxycholate, 0.1% SDS 1% NP-40) containing complete protease inhibitor cocktail (1x; Roche). Protein concentrations were determined by BCA Protein Assay Kit (Pierce) and equal amounts of proteins or equal volumes of iAsCM were subjected to standard SDS-PAGE, followed by transfer to PVDF membranes. No-Stain Protein Labelling Reagent (Invitrogen) was used to confirm equal transfer between samples. Membranes were blocked for 45 min with 5% milk powder in TBS containing 0.5% Tween-20 (TBST) and incubated primary antibodies (Tab. 4) overnight at 4°C, followed by incubation with horse radish peroxidase-conjugated secondary antibody (diluted 1:5000) for 90 min at room temperature. Immunoreactive bands in cell extracts were visualized using ECL Western Blotting Detection Reagent (Amersham) on iBright CL1500 Imaging System (Invitrogen). For murine synaptosomal extracts, immunoreactive signals were visualized using SuperSignal West Femto Maximum Sensitivity Substrate (Thermo Scientific). Densitometric analyses were performed using Fiji software.

### Immunocytochemistry

For immunocytochemical analysis, cells were plated on Matrigel-coated 13 mm-diameter glass slides in 24-well plates. For analysis, the cells were washed with PBS and fixed in 4% PFA in PBS for 20 min, followed by in PBS-based blocking/permeabilization solution containing 0.1% Triton X-100 and 5% donkey serum for 1 hour at room temperature. Primary antibody dilutions (Tab. 4) were applied overnight at 4°C in PBS. Thereafter, the cells were washed twice in PBS and once in PBST, followed by incubation with Alexa Fluor-conjugated secondary antibodies for 2 hours at room temperature (Tab. 4). For nuclear staining, DAPI (Thermo Fisher) diluted in PBS was used. All images were acquired on Zeiss LSM780 or Zeiss LSM800 confocal microscopes and analyzed using Fiji software.

### Analysis of gene transcription

Total RNA was extracted from tissue or cells using RNeasy Mini/Micro Kit (Qiagen), treated with RNase-Free DNase Set (Qiagen), and reverse transcribed using the High Capacity RNA-to-cDNA Kit (Applied Biosystems) according to the manufacturers’ protocols. Quantitative PCR were carried out using the TaqMan Fast Advanced Master Mix (Applied Biosystems) and TaqMan assays using QuantStudio 7 Flex Real-Time PCR System (Applied Biosystems) or Quant Studio 6 Pro Real-Time PCR System (Applied Biosystems). TaqMan assay details are listed in Tab. 5. The fold change in transcript levels was calculated using the cycle threshold (CT) comparative method (2^−ddCT^) normalizing to CT values of the internal control gene *GAPDH/Gapdh.* Samples were run in triplicates and non-template reaction served as negative controls.

To assess the pluripotency of iPSC line, the TaqMan Scorecard Assay (Applied Biosystems) was used according to the manufacturer’s instructions. RNA extraction and cDNA reverse transcription of 1 µg of total RNA was performed as described above. The scorecard assay was run in a 384-well format on QuantStudio 7 Flex Real-Time PCR System using the Standard Curve Method. Gene expression data were analyzed using the web-based hPSC Scorecard Analysis Software (Thermo Fisher Scientific).

The expression of 84 PPAR target genes was assessed using the Human PPAR Targets RT^2^ Profiler PCR Array (Qiagen; 384-well format) according to the manufacturer’s instructions. RNA was extracted using RNeasy Mini/Micro Kit (Qiagen), treated with RNase-Free DNase Set (Qiagen), and reverse transcribed with RT^2^ First Strand Kit (Qiagen). Quantitative PCR was performed using RT^2^ SYBR Green ROX qPCR Mastermix (Qiagen) on a QuantStudio 7 Flex Real-Time PCR System (Applied Biosystems). The comparison of transcript levels was calculated using the cycle threshold (CT) comparative method (2^−dCT^) normalizing to CT values of internal control gene *GAPDH*. Data analysis was conducted with the web-based GeneGlobe Data Analysis Center (https://geneglobe.qiagen.com). Changes in mRNA levels in E3KO and E4 KO cells were assessed as compared to the respective isogenic WT controls. Genes with more than 2-fold difference in expression were considered to differentially expressed. Two arrays were run for each E3WT vs E3KO and E4WT vs E4KO comparison and genes showing the same changes in both arrays were chosen for further validation with TaqMan qRT-PCR assays.

### Analysis of mitochondria

For quantification of the amounts of mitochondria in iNs, the cells were cultured on Matrigel-coated 13 mm-diameter glass slides in 24-well plates and incubated with 100 nM MitoTracker Red CMXRos (Invitrogen) for 30 min. After staining, cells were washed with fresh medium and fixed in 4% PFA in PBS for 20 min. Images were acquired using a Zeiss LSM800 confocal microscope. Z-stack images (14 slices; interval 0.2 μm) were analyzed using Fiji software. To measure cell body area, GFP signal was used to manually track each cell body shape. Images of mitochondria were binarized and thresholding parameters adjusted based on the MitoTracker signal. Th particle analysis function was used to measure mitochondria area obtained by pixels that make up the MitoTracker signal. The fraction of the total GFP+ cell area occupied by MitoTracker Red+ mitochondria was obtained by dividing total mitochondria area by cell area.

### *APOE* genotyping

Genomic DNA was extracted from stem cells using Wizard Genomic DNA Purification Kit (Promega). *APOE* genotyping was carried out using the TaqMan SNP Genotyping Assay and TaqMan Genotyping Master Mix (Applied Biosystems) on a QuantStudio 7 Flex Real-Time PCR System (Applied Biosystems). Genotypes were determined based on single nucleotide polymorphism rs429358 that defines ε3 and ε4 alleles of human *APOE* (Assay ID C_3084793_20; Applied Biosystem). FAM and VIC reporter dyes were used for allele discrimination.

### Statistical analysis

Statistical analyses were performed using GraphPad Prism. The applied statistical tests and the number *n* (sample size) are indicated in each figure legends. Data are presented as the mean ± SD, unless otherwise stated in the respective figure legends.

## METHOD TABLES

**Table 1:** Ultra-high performance liquid chromatography (UHPLC) gradient for LC part. A: water with 0.2% formic acid; B: acetonitrile with 0.2% formic acid.

| No | LC1 |  |  |  | LC2 |  |  |  |
| --- | --- | --- | --- | --- | --- | --- | --- | --- |
|  | Time (min) | Flow (ml/min) | A (%) | B (%) | Time (min) | Flow (ml/min) | A (%) | B (%) |
| 1 | 0.00 | 0.8 | 100 | 0 | 0.00 | 0.8 | 100 | 0 |
| 2 | 0.25 | 0.8 | 100 | 0 | 0.25 | 0.8 | 100 | 0 |
| 3 | 1.50 | 0.8 | 88 | 12 | 0.50 | 0.8 | 75 | 25 |
| 4 | 2.70 | 0.8 | 82.5 | 17.5 | 2.00 | 0.8 | 50 | 50 |
| 5 | 4.00 | 0.8 | 50 | 50 | 3.00 | 0.8 | 25 | 75 |
| 6 | 4.50 | 0.8 | 0 | 100 | 3.50 | 0.8 | 0 | 100 |
| 7 | 4.70 | 1.0 | 0 | 100 | 4.70 | 1.0 | 0 | 100 |
| 8 | 5.00 | 1.0 | 0 | 100 | 5.00 | 1.0 | 0 | 100 |
| 9 | 5.10 | 1.0 | 100 | 0 | 5.10 | 1.0 | 100 | 0 |
| 10 | 5.80 | 1.0 | 100 | 0 | 5.80 | 1.0 | 100 | 0 |

**Table 2:** Ultra-high performance liquid chromatography (UHPLC) gradient for flow injection. B: FIA solution in methanol.

| No | Time (min) | Flow (ml/min) | A (%) | B (%) |
| --- | --- | --- | --- | --- |
| 1 | 0.0 | 0.03 | 0 | 100 |
| 2 | 1.6 | 0.03 | 0 | 100 |
| 3 | 2.4 | 0.20 | 0 | 100 |
| 4 | 2.8 | 0.20 | 0 | 100 |
| 5 | 3.0 | 0.03 | 0 | 100 |

**Table 3:** Acquisition method parameters for liquid chromatography (LC) and flow injection analysis (FIA). n/a, not applicable; MS, mass spectrometry; MRM, multiple reaction monitoring.

| Option | Parameter | LC1 | LC2 | FIA1 | FIA2 |
| --- | --- | --- | --- | --- | --- |
| **MS** | Scan type | MRM | MRM | MRM | MRM |
|  | Polarity | positive | negative | positive | positive |
|  | MRM detection window (sec) | 30 | 30 | - | - |
|  | Target scan time (sec) | 0.15 | 0.15 | - | - |
|  | Duration (min) | 5.45 | 5.45 | 2.95 | 2.95 |
|  | Delay time (sec) | 0 | 0 | 0 | 0 |
|  | Cycle (sec) | 0.25 | 0.15 | n/a | n/a |
| **Advanced MS** | Resolution Q1 | unit | unit | unit | unit |
|  | Resolution Q3 | unit | unit | unit | unit |
|  | Intensity threshold | 0 | 0 | 0 | 0 |
|  | Setting time (ms) | 0 | 0 | 0 | 0 |
|  | Pause between mass ranges (ms) | 2 | 2 | 5.007 | 3 |
| **Source/ gas** | Curtain gas | 45 | 20 | 20 | 10 |
|  | Collision gas | 9 | 8 | 9 | 9 |
|  | Ion spray voltage | 5500 | -4500 | 5500 | 5500 |
|  | Temperature | 500 | 650 | 200 | 350 |
|  | Ion source gas 1 | 60 | 40 | 40 | 30 |
|  | Ion source gas 2 | 70 | 40 | 50 | 85 |

**Table 4:**
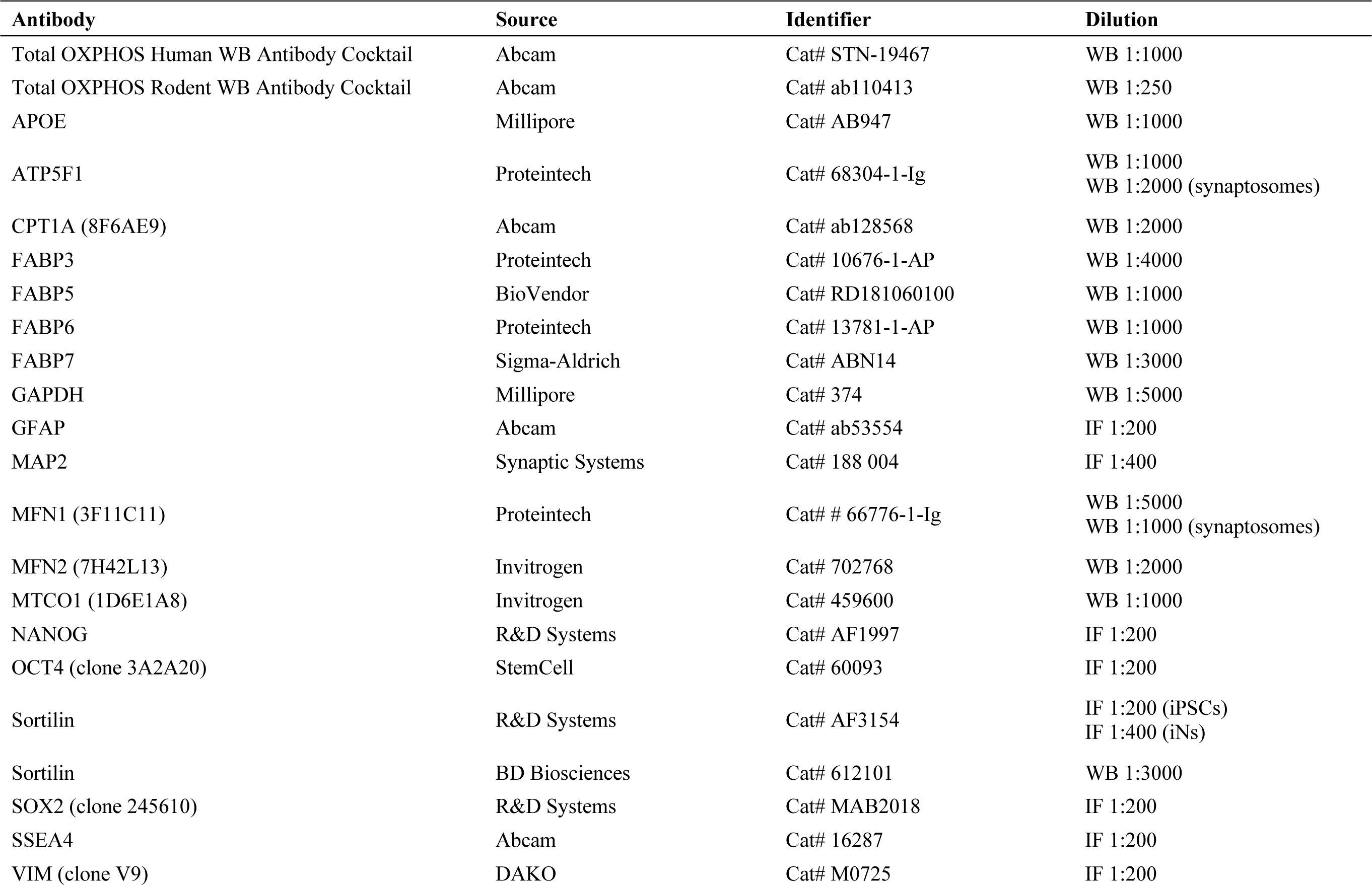
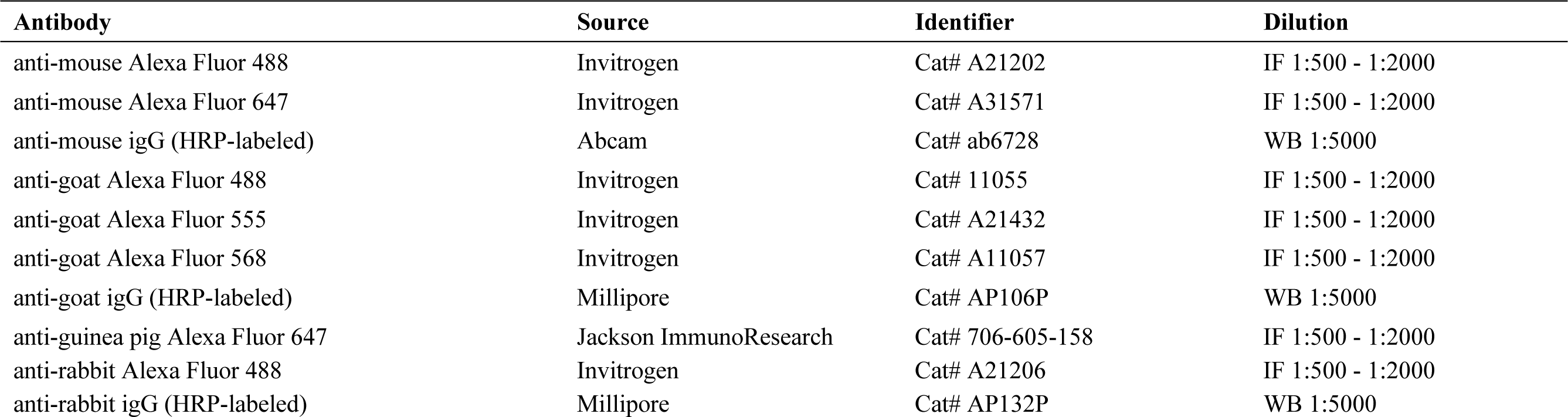
Primary and secondary antibodies used in this study.

**Table 5:**
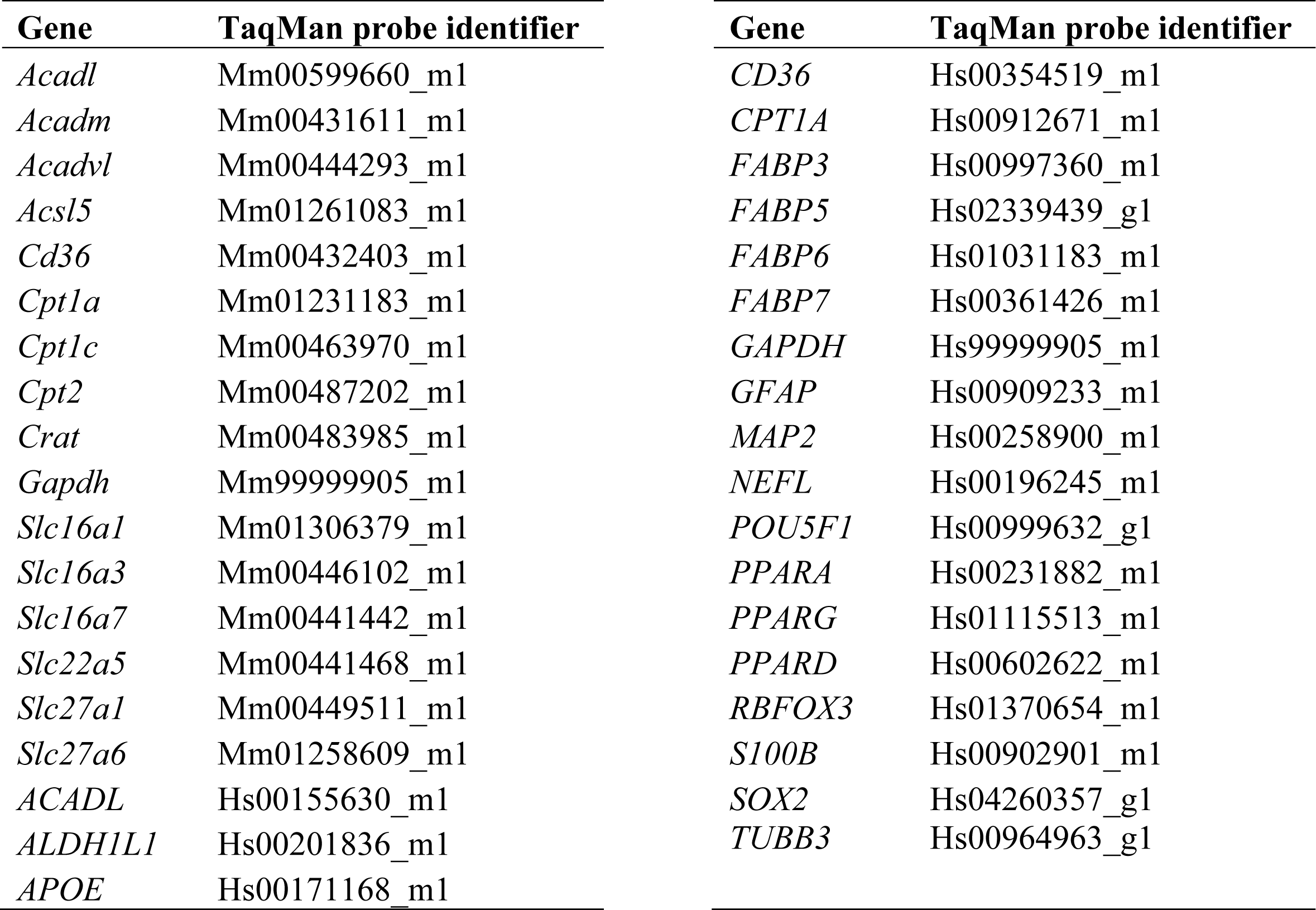
Taqman probes used to determine transcript levels of murine and human genes.

