## Supplementary figures for "Interaction of sortilin with apolipoprotein E3 enables neurons to use long-chain fatty acids as alternative metabolic fuel"

### SUPPLEMENTARY FIGURES AND FIGURE LEGENDS

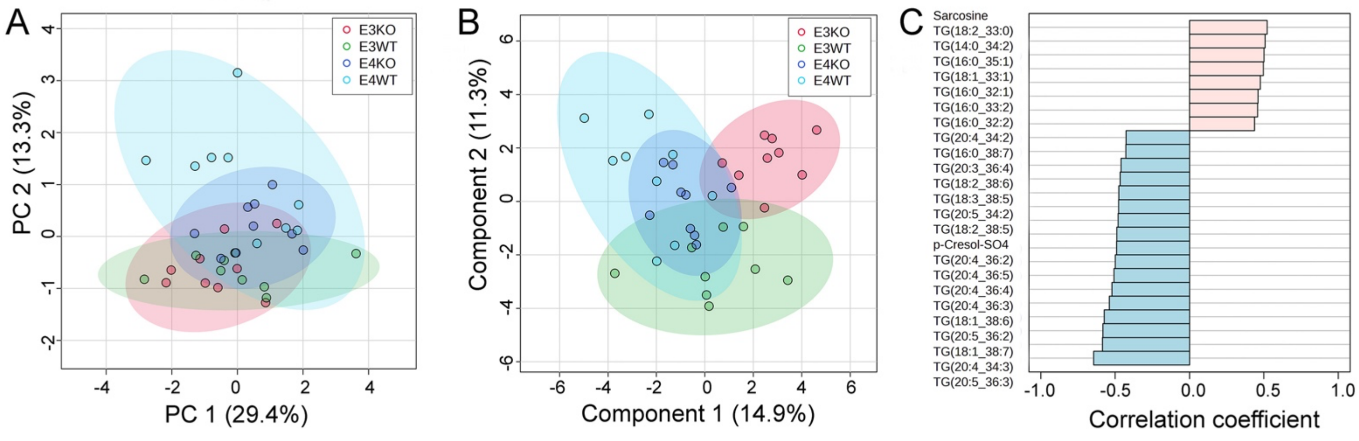


**Supplementary figure 1 (related to figure 1): Global metabolomics of mouse plasma samples**

(**A, B**) EDTA-treated plasma samples from male mice of the indicated *Sort1* and human *APOE* genotypes (age 12 weeks, n=10 per genotype group) were subjected targeted metabolomics using LC-MS and FIA-MS analysis (MxP Quant 500 kit, Biocrates Life Science). Data were analyzed using MetaboAnalyst 5.0 (https://www.metaboanalyst.ca/). Unsupervised Principal Component Analysis (A) and supervised Partial Least-Squares Discriminant Analysis (sparsed variant) (B) were used to query distribution of samples according to genotype (E3WT, green; E3KO, red; E4WT, light blue; E4KO, dark blue). The performance of PLS-DA model was evaluated using 5-fold cross validation (classification error rate was 39% for 2 components). (**C**) Pattern hunter analysis of the plasma metabolome data set identifying metabolites characteristic of sortilin and apoE3 interaction with E3WT being different to the other three genotype groups. The X-axis gives the calculated correlation coefficients, reflecting positive (pink) or negative (blue) correlation of the respective metabolite concentration with the tested pattern.

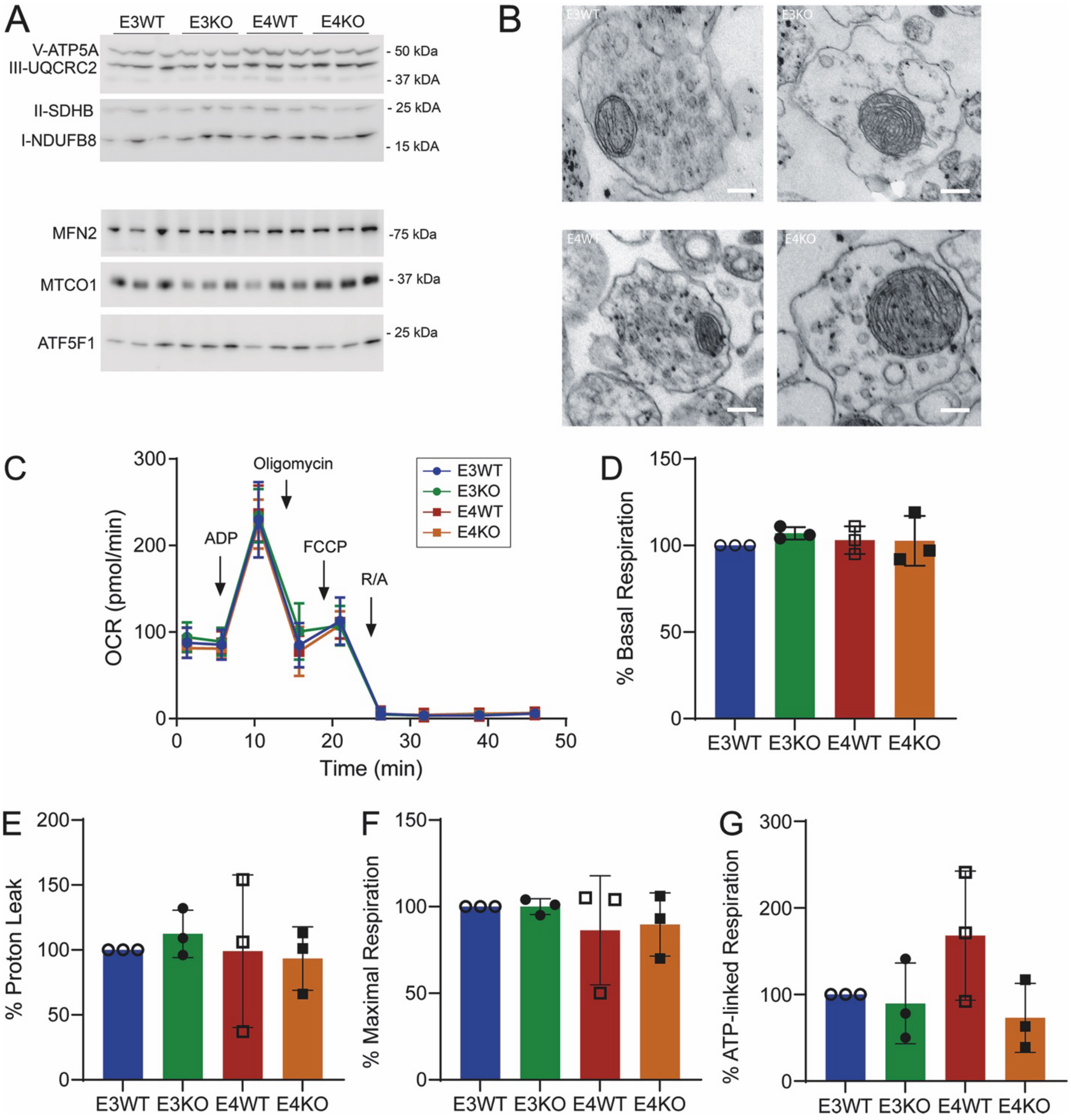


**Supplementary figure 2 (related to figure 2): Analysis of murine mitochondria structure and liver function**

(**A**) Western blot analysis of respiratory chain subunits and mitofusin 2 (MFN2) in synaptosomes isolated from E3WT, E3KO, E4WT, or E4KO mouse brains (n=3 animals per group). Five µg synaptosomal lysate were loaded in each lane. (**B**) Representative electron microscopic images of synaptosomes isolated from E3WT, E3KO, E4WT, or E4KO mouse brain cortices showing trapped neuronal mitochondria. Scale bars: 100 nm. (**C**) Respiration profiles of mitochondria isolated from livers of male mice of the indicated genotypes (12 weeks of age). Oxidation consumption rates (OCR) was measured by sequential addition of ADP, oligomycin, carbonyl cyanide-p trifluoromethoxyphenylhydrazone (FCCP), as well as rotenone and antimycin A (R/A) at the indicated time points. (**D - G**) Quantification of basal respiration (D), proton leak (E), maximal OCR (F), and ATP-linked respiration (G) in liver mitochondria of the indicated genotypes. Data are given as % of the E3WT condition (set to 100%). Data represent the mean ± SD from n=3 animals per genotype with n=20-24 technical replicates per data point. Statistical significance of data was tested using 2way ANOVA with Tukey's multiple comparison test (*, $p < 0.05$; **, $p < 0.01$; ***, $p < 0.001$; ****, $p < 0.0001$).

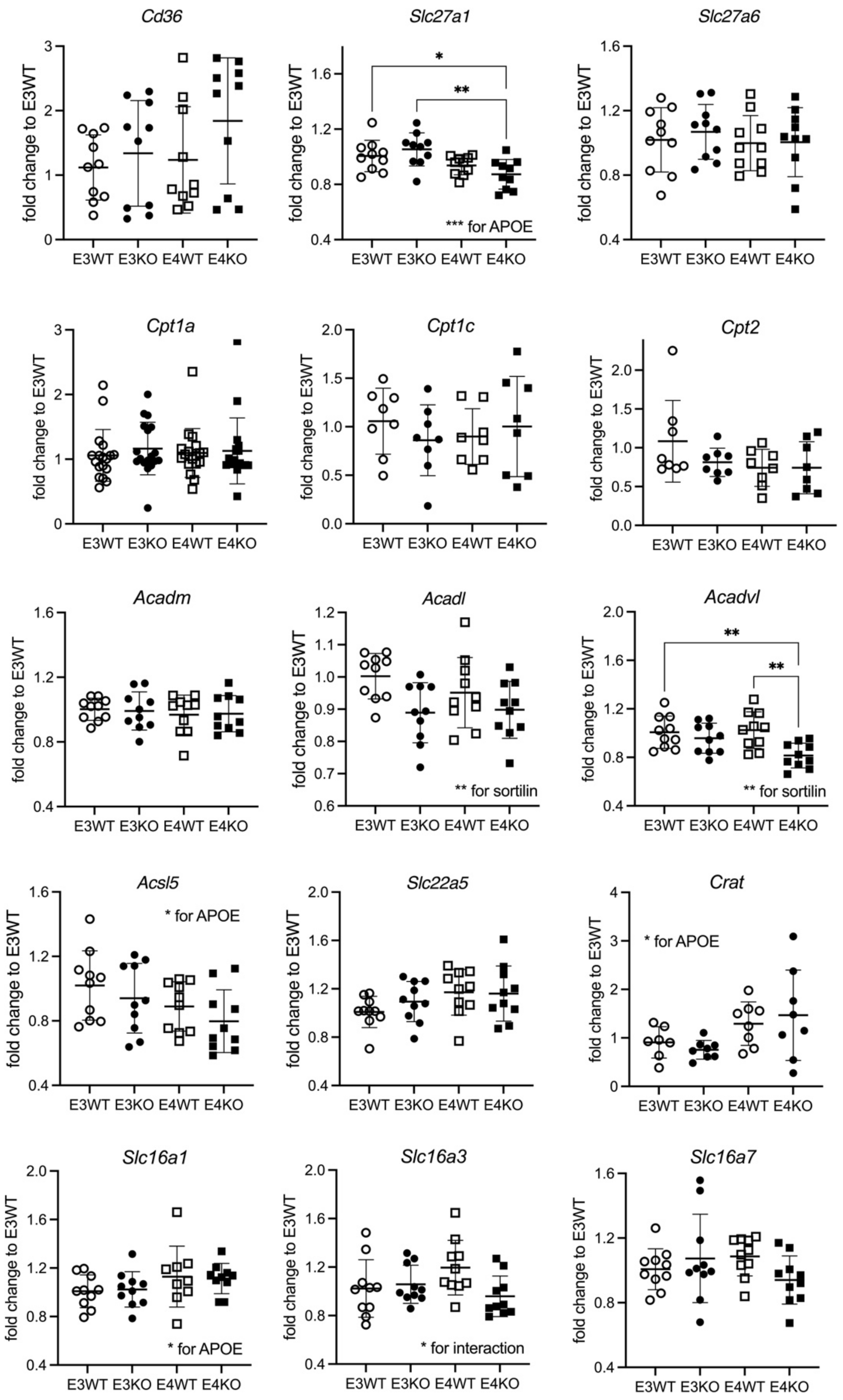


**Supplementary figure S3 (related to figure 3): Gene expression patterns in murine brain cortices related to mitochondrial consumption of long-chain fatty acids**

Cortical gene expression patterns were studied in male mice (12 weeks of age) of the indicated *Sort1* and human *APOE* genotypes using quantitative RT-PCR (n=10-18 animals per group). Individual data points as well as the mean ± SD of the entire genotype group are given. Statistical significance of data was tested using 2way ANOVA with Tukey's multiple comparison test (*, $p < 0.05$; **, $p < 0.01$; ***, $p < 0.001$). *Acadl*, acyl-CoA dehydrogenase long chain; *Acadm*, acyl-CoA dehydrogenase medium chain; *Acadvl*, acyl-CoA dehydrogenase very long chain; *Acsl5*, acyl-CoA synthetase long chain family member 5; *Cd36*, fatty acid translocase; *Cpt1a*, carnitine O-palmitoyltransferase 1A; *Cpt1c*, carnitine O-palmitoyltransferase 1C; *Cpt2*, carnitine O-palmitoyltransferase 2; *Crat*, carnitine O-acyltransferase; *Slc16a1*, solute carrier family 16 member 1 encoding monocarboxylate transporter (MCT) 1; *Slc16a3*, solute carrier family 16 member 3 encoding MCT4; *Slc16a7*, solute carrier family 16 member 7 encoding MCT2; *Slc22a5*, solute carrier family 22 member 5; *Slc27a1*, solute carrier family 27 member 1 encoding fatty acid transport protein 1 (FATP1); *Slc27a6*, solute carrier family 27 member 6, encoding FATP6.

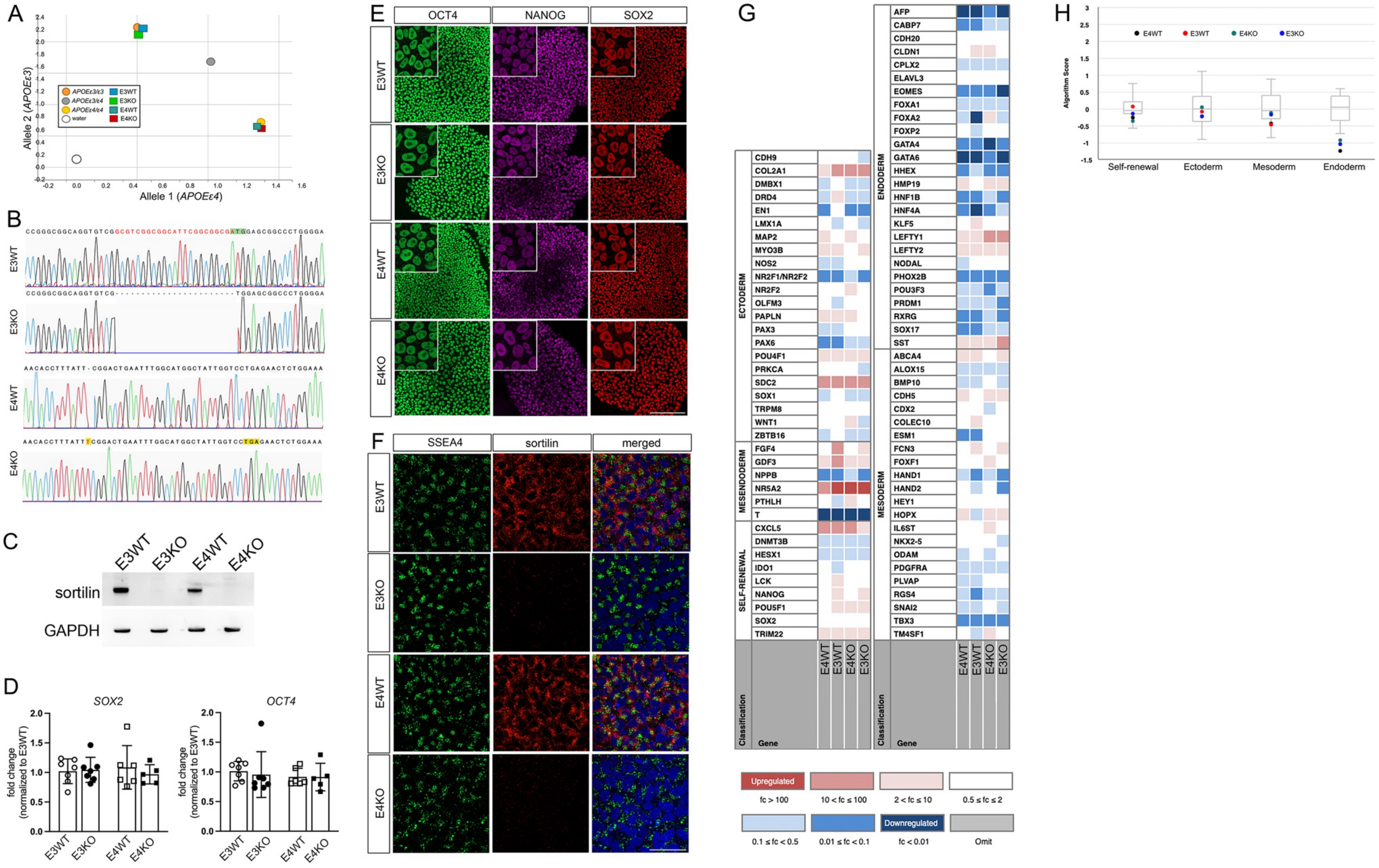
A
Allele 2 (APOEε3)
Allele 1 (APOEε4)
APOEε3/ε3
APOEε3/ε4
APOEε4/ε4
water
E3WT
E3KO
E4WT
E4KO
B
E3WT
E3KO
E4WT
E4KO
CCGGGCGGCAGGTGTCGGCGTCGGCGGCATTCGGCGGCGATGGAGCGGCCCTGGGGA
CCGGGCGGCAGGTGTCG--------------------TGGAGCGGCCCTGGGGA
AACACCTTTATT-CGGACTGAATTTGGCATGGCTATTGGTCCTGAGAACTCTGGAAA
AACACCTTTATTTCGGACTGAATTTGGCATGGCTATTGGTCCTGAGAACTCTGGAAA
C
sortilin
GAPDH
D
SOX2
OCT4
fold change (normalized to E3WT)
E
OCT4
NANOG
SOX2
F
SSEA4
sortilin
merged
G
ECTODERM
MESENDODERM
SELF-RENEWAL
ENDODERM
MESODERM
Classification
Gene
CDH9
COL2A1
DMBX1
DRD4
EN1
LMX1A
MAP2
MYO3B
NOS2
NR2F1/NR2F2
NR2F2
OLFM3
PAPLN
PAX3
PAX6
POU4F1
PRKCA
SDC2
SOX1
TRPM8
WNT1
ZBTB16
FGF4
GDF3
NPPB
NR5A2
PTHLH
T
CXCL5
DNMT3B
HESX1
IDO1
LCK
NANOG
POU5F1
SOX2
TRIM22
AFP
CABP7
CDH20
CLDN1
CPLX2
ELAVL3
EOMES
FOXA1
FOXA2
FOXP2
GATA4
GATA6
HHEX
HMP19
HNF1B
HNF4A
KLF5
LEFTY1
LEFTY2
NODAL
PHOX2B
POU3F3
PRDM1
RXRG
SOX17
SST
ABCA4
ALOX15
BMP10
CDH5
CDX2
COLEC10
ESM1
FCN3
FOXF1
HAND1
HAND2
HEY1
HOPX
IL6ST
NKX2-5
ODAM
PDGFRA
PLVAP
RGS4
SNAI2
TBX3
TM4SF1
Upregulated
fc > 100
10 < fc ≤ 100
2 < fc ≤ 10
0.5 ≤ fc ≤ 2
0.1 ≤ fc < 0.5
0.01 ≤ fc < 0.1
Downregulated
fc < 0.01
Omit
H
Algorithm Score
Self-renewal
Ectoderm
Mesoderm
Endoderm

**Supplementary figure S4 (related to figure 4): Generation of human induced pluripotent stem cells lines with defined *APOE* and *SORT1* genotypes**

**(A)** *APOE* genotyping of human iPS cell (hiPSC) lines using quantitative RT-PCR. The scatter plot X- and Y-axes represent allele discrimination for *APOEε3* and *APOEε4* genotypes. Shown hiPSC lines (colored squares) are *APOEε3* (E3) or *APOEε4* (E4) and either wildtype (WT) or genetically deficient for *SORT1* (KO). As internal controls, human samples being *APOEε3/ε3* (orange dot), *APOEε3/ε4* (grey dot), or *APOEε4/ε4* (yellow dot) were included in the genotyping PCR. The white dot represents the water-only control. **(B)** Genome sequence analysis of the *SORT1* gene region in genome edited iPSC lines. Cell line E3KO carries a 23 base pair deletion that includes the ATG (green), as compared to the isogenic E3WT control line. Cell line E4KO harbors a one nucleotide insertion (red), resulting in premature stop codon (TGA, yellow), as compared to the isogenic E4WT control. Sequences were aligned to the human *SORT1* reference sequence (NCBI: BC023542.1). **(C)** Western blot analyses of lysates from the indicated iPSC lines, documenting the presence of sortilin in E3WT and E4WT, but absence of protein from lines E3KO and E4KO. Detection of GAPDH served as loading control. **(D)** Transcript levels of *SOX2* and *OCT4* in iPSC lines of the indicated *SORT1* and *APOE* genotypes were tested using qRT-PCR. Data points for n=5-7 independent cell cultures as well as mean ± SD of the entire data set are given. **(E, F)** Representative immunofluorescence images of iPSC lines of the indicated genotypes stained for pluripotency markers OCT4 (green), NANOG (purple), and SOX2 (red) (in E) as well as for SSEA4 (green) and sortilin (red). In F, DAPI (blue) in merge. Scale bars: 200 µm (E), 50 µm (F). **(G)** Transcript levels of 94 genes involved in pluripotency and tri-lineage differentiation potential were quantified in iPSC lines of the indicated *SORT1* and *APOE* genotypes using the TaqMan hPSC Scorecard Assay (as detailed in STAR method section). Expression levels are shown as heatmap with colors correlating to the fold change in expression of the indicated gene relative to the undifferentiated reference set. **(H)** Box plot depicting transcript levels of genes in the Scorecard Assay panel associated with self-renewal or ectodermal, mesodermal or endodermal differentiation in iPSC lines of the indicated genotypes. Sample scores are plotted in color. Gray box and whisker plots represent undifferentiated reference datasets.

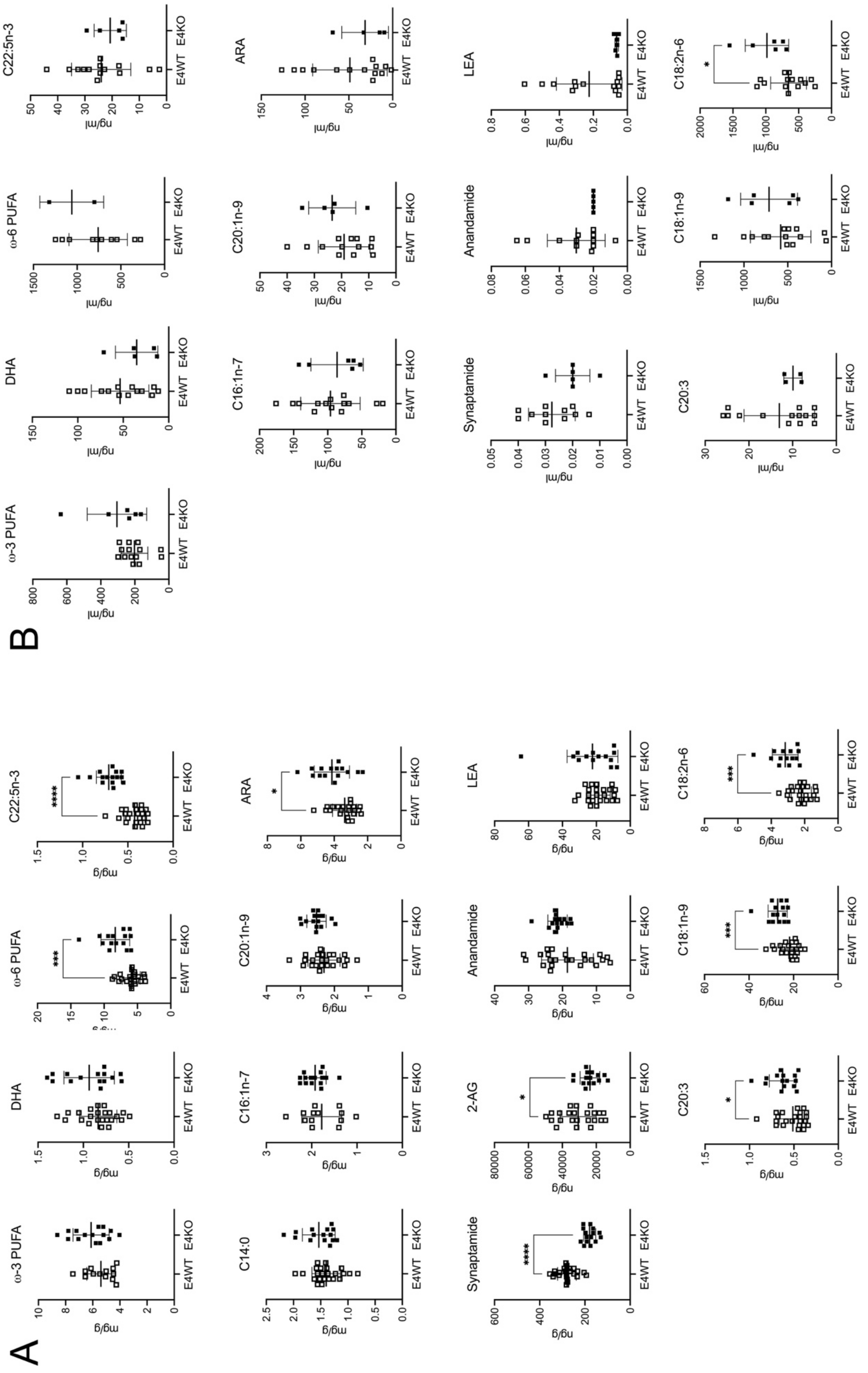
A
ω-3 PUFA
DHA
ω-6 PUFA
C22:5n-3
C14:0
C16:1n-7
C20:1n-9
ARA
Synaptamide
2-AG
Anandamide
LEA
C20:3
C18:1n-9
C18:2n-6
B
ω-3 PUFA
DHA
ω-6 PUFA
C22:5n-3
C16:1n-7
C20:1n-9
ARA
Synaptamide
Anandamide
LEA
C20:3
C18:1n-9
C18:2n-6
E4WT
E4KO
mg/g
ng/g
ng/ml

**Supplementary figure S5 (related to figure 5): Sortilin deficiency does not impact the metabolism of poly-unsaturated fatty acids in human apoE4 neurons**

(**A**) Concentrations of selected lipids in iPSC-derived cortical neurons were determined using LC-MS. Neurons (day 12 of culture) were *APOEε4/ε4* and either wildtype (E4WT) or genetically deficient for *SORT1* (E4KO). Data from individual biological replicates (n=15-23) of 2 - 4 individual differentiation experiments are given. (**B**) Concentrations of selected lipids in the cell supernatant of iPSC-derived E4WT and E4KO astrocytes (days 21- 28 of culture) were determined using LC-MS. Individual biological replicates (n=6-14) of 2 - 6 individual differentiation experiments are shown. Individual data points as well as mean ± SD of the entire genotype group are given. Statistical significance of data was tested using unpaired Student's *t* test (two-tailed; *, $p < 0.05$; **, $p < 0.01$; ***, $p < 0.001$; ****, $p < 0.0001$. ARA, arachidonic acid; 2-AG, 2-arachidonoylglycerol; DHA, docosahexaenoic acid; LEA, linoleoyl ethanolamide; PUFA, poly-unsaturated fatty acid.

A

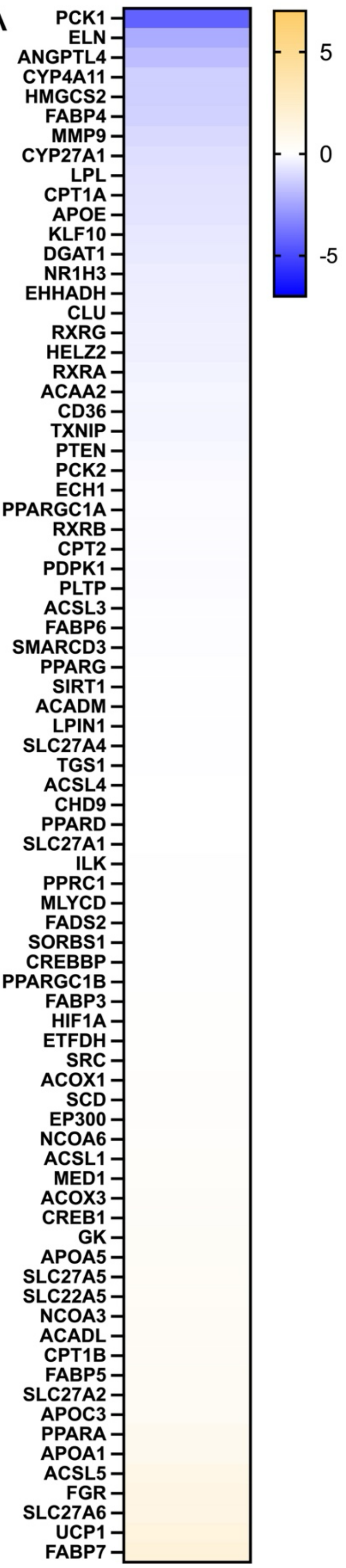


B

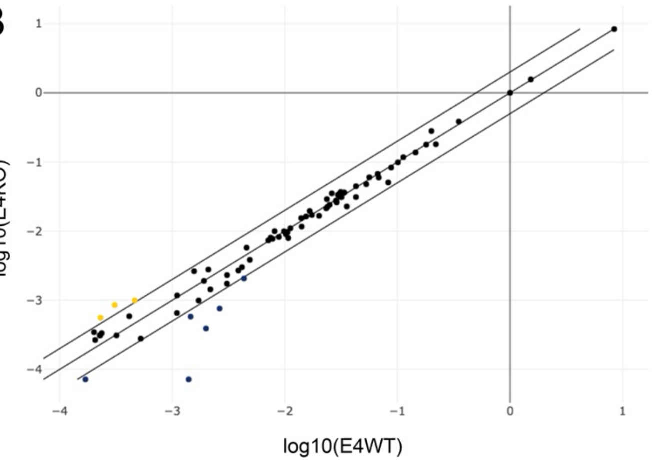


C

**Genes Under-Expressed in**
**E4KO vs. Control Group (E4WT)**

| Gene Symbol | Fold Regulation | |
|---|---|---|
| | #1 | #2 |
| ANGPTL4 | -3.47 | -5.32 |
| ELN | -5.10 | -2.84 |
| FABP4 | -2.36 | -3.99 |
| MMP9 | -2.10 | -5.75 |

D

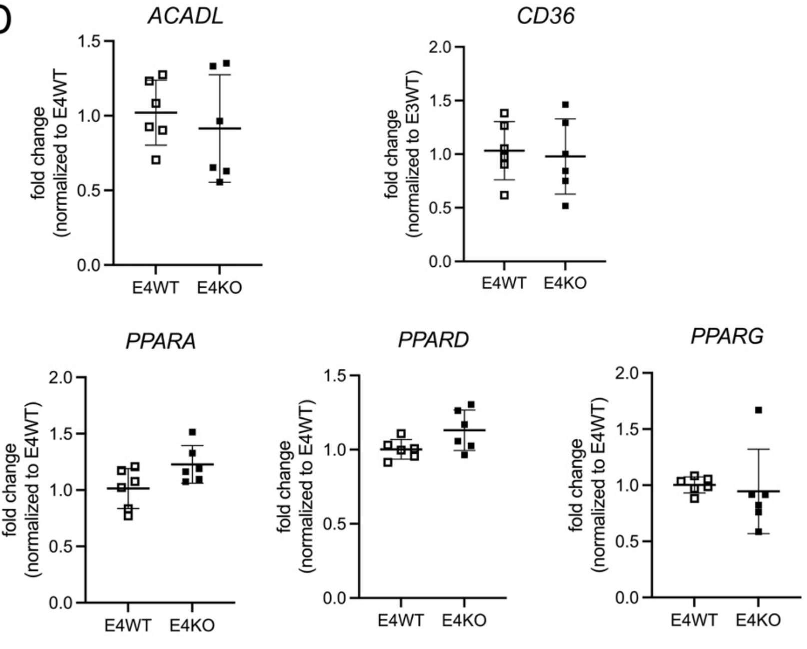

**Supplementary figure S6 (related to figure 6): Sortilin deficiency does not impair PPAR-dependent gene expression in human apoE4 neurons**

**(A - C)** Expression levels of 84 peroxisome proliferator-activated receptor (PPAR) target genes were determined in iPSC-derived human neurons treated with astrocyte-conditioned media using the human PPAR Targets $RT^2$ Profiler PCR Array. Neurons (day 12 of culture) were *APOEε4/ε4* and either wildtype (E4WT) or genetically deficient for *SORT1* (E4KO). One exemplary experiment of n=2 biological replicates is shown. (A) Expression levels are given as heat map with transcript levels in E4KO presented as $\log_2$ fold change as compared to E4WT (set to 0). Blue and yellow color spectra indicate down-regulated or up-regulated genes, respectively. (B) Scatterplot analysis comparing log-transformed relative expression for all tested PPAR target genes in E4WT and E4KO neurons. The center diagonal line indicates unchanged gene expression levels. The outer diagonal lines indicate 2-fold regulation threshold. Genes with >2-fold difference in transcript level between groups are presented as blue and yellow dots, indicating down- or up-regulated genes, respectively. **(C)** Selected list of genes downregulated by >2 fold in E4KO as compared to E4WT human neurons treated with iAs-conditioned media. Data from two independent experiments are shown. **(D)** Quantitative RT-PCR analysis of selected PPAR target genes in E4WT and E4KO human neurons treated with iAs-conditioned media (day 12 of culture). Data points for n=6 biological replicates (from 2 differentiations) as well as mean ± SD are given. No statistical difference comparing genotypes was seen using unpaired Student's *t* test (two-tailed).

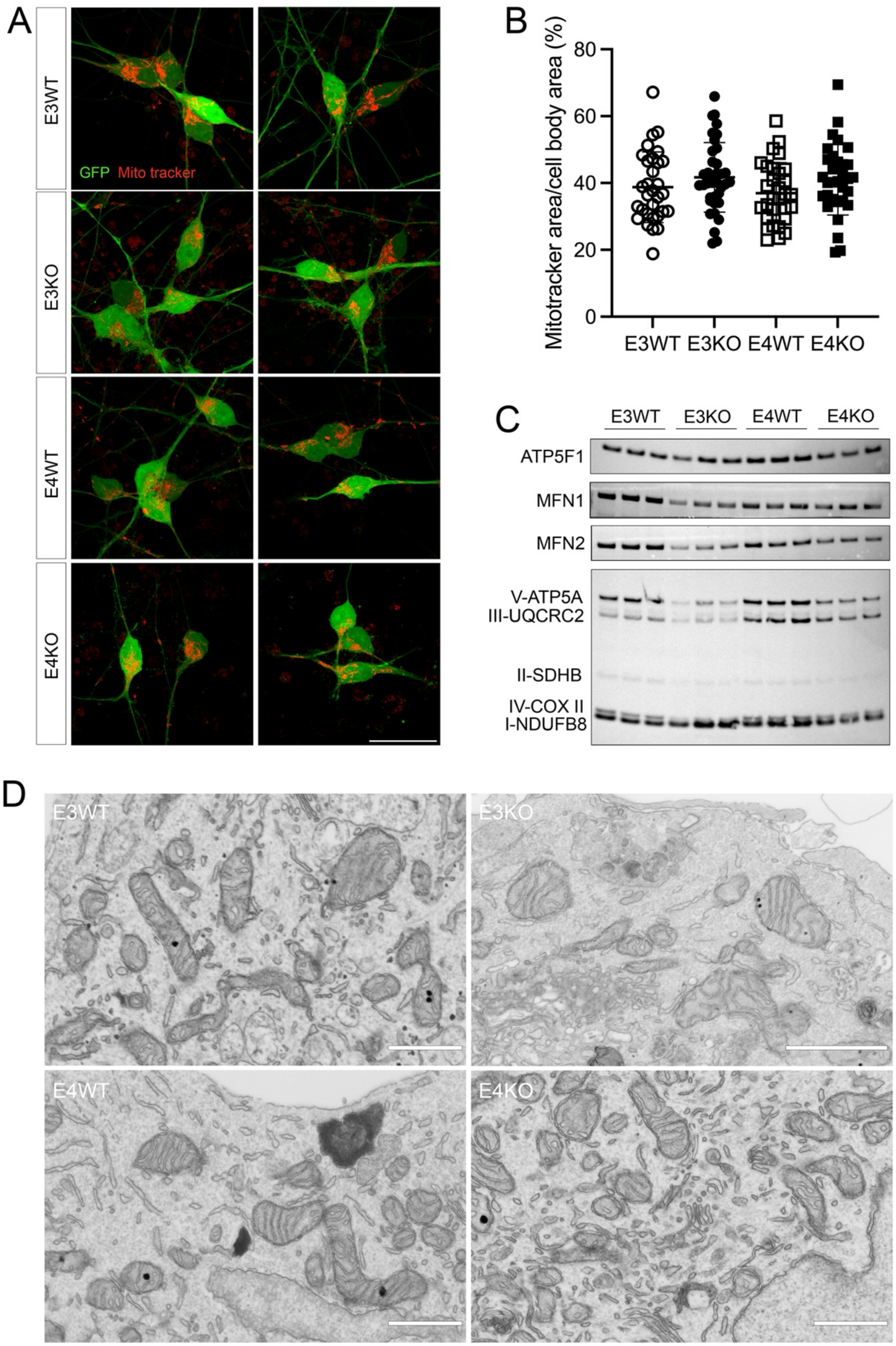


**Supplementary figure S7 (related to figure 7): Loss of sortilin or the presence of apoE4 does not impact mitochondrial appearance in human iNs.**

**(A)** Representative fluorescence images of human neurons of the indicated genotypes (day 7 of culture) depicting native fluorescence for GFP (green) as well as MitoTracker Red CMXRos (red). Scale bar: 25 µm. **(B)** Quantification of MitoTracker Red CMXRos-positive cell area (expressed as % of the total soma area) in human neurons. Data points represent individual cell measurements (n=30-37) from 2 differentiation experiments. Individual data points as well as mean ± SD of the entire genotype group are shown. **(C)** Levels of the indicated respiratory chain subunits and mitofusins (MFN1 and MFN2) were determined in 5 µg of total lysates from human neurons of the indicated *SORT1* and *APOE* genotypes using Western blotting (day 7 of culture, 2 individual differentiation experiments). **(D)** Representative electron microscopic images of mitochondria in human neurons of the indicated *SORT1* and *APOE* genotypes. Scale bars: 730 nm or 1 µm (E3KO).
